# Quality assessment and control of unprocessed anatomical, functional, and diffusion MRI of the human brain using *MRIQC*

**DOI:** 10.1101/2024.10.21.619532

**Authors:** McKenzie P. Hagen, Céline Provins, Eilidh MacNicol, Jamie K. Li, Teresa Gomez, Mélanie Garcia, Saren H. Seeley, Jon Haitz Legarreta, Martin Norgaard, Patrick G. Bissett, Russell A. Poldrack, Ariel Rokem, Oscar Esteban

## Abstract

Quality control of MRI data prior to preprocessing is fundamental, as substandard data are known to increase variability spuriously. Currently, no automated or manual method reliably identifies subpar images, given pre-specified exclusion criteria. In this work, we propose a protocol describing how to carry out the visual assessment of T1-weighted, T2-weighted, functional, and diffusion MRI scans of the human brain with the visual reports generated by *MRIQC*. The protocol describes how to execute the software on all the images of the input dataset using typical research settings (i.e., a high-performance computing cluster). We then describe how to screen the visual reports generated with *MRIQC* to identify artifacts and potential quality issues and annotate the latter with the “rating widget” – a utility that enables rapid annotation and minimizes bookkeeping errors. Integrating proper quality control checks on the unprocessed data is fundamental to producing reliable statistical results and crucial to identifying faults in the scanning settings, preempting the acquisition of large datasets with persistent artifacts that should have been addressed as they emerged.

**RELATED LINKS:** *Key reference(s) using this protocol:* Esteban, O. et al. (2017), PLoS ONE 12(9): e0184661. [10.1371/journal.pone.0184661] Esteban, O. et al. (2019), Sci Data 6, 30. [10.1038/s41597-019-0035-4] Esteban, O. et al. (2020), Nat Prot 15, 2186–2202. [10.1038/s41596-020-0327-3] Provins, C. et al. (2023), Front. Neuroinform. 1, 2813-1193. [10.3389/fnimg.2022.1073734] Bissett P. et al. (2024) Sci Data 11: 809. [10.1038/s41597-024-03636-y]

*Key data used in this protocol:* Amsterdam Open MRI Collection: Population Imaging of Psychology^1^ (AOMIC-PIOP1; ds002785 [https://openneuro.org/datasets/ds002785]).

## Introduction

Ensuring the quality of acquired data is a crucial initial step for any magnetic resonance imaging (MRI) analysis workflow because low-quality images may alter the statistical outcomes^2–5^. Therefore, a quality assurance (QA) check on the reconstructed images as they are produced by the scanner – i.e., “*acquired*” or “*unprocessed*” data– is fundamental to identify subpar instances and prevent their progression through analysis when they meet predefined exclusion criteria. Data curation protocols^6^ for MRI data are still being established, and automated tools to carry out quality control (QC) are still in their infancy ^7,8^. Laboratories currently address the challenge of QC by applying their internal knowledge to visual assessments or by accounting for artifacts and other quality issues in the subsequent statistical analysis. One common practice for doing QC of unprocessed data entails screening every slice of every scan individually^9^, which is time-consuming, subjective, prone to human errors, and variable within and between raters. Additionally, QA/QC protocols may vary substantially across MRI modalities due to available visualization software or knowledge about modality-specific artifacts. These unstandardized protocols can result in low intra- and inter-rater reliability, hindering the definition of objective exclusion criteria in studies. Intra-rater variability derives from aspects such as training, subjectivity, varying annotation settings and protocols, fatigue, or bookkeeping errors. The difficulty in calibrating between experts and the lack of agreed exclusion criteria, which are contingent on each particular application^10^, lies at the heart of inter-rater variability. Adhering to a well-developed standard operating procedure (SOP) that describes the QC process can minimize these variabilities.

With the current data deluge in neuroimaging, manual QC of every scan has become onerous for typically-sized datasets and impractical for consortium-sized datasets, exacerbating the problems mentioned above. For the smaller datasets, QC can be streamlined by using informative visualizations to rate images and minimize bookkeeping effort. For large-scale databases, early approaches^11–14^ to objectively estimate image quality have employed “image quality metrics” (IQMs) that quantify objectively defined, although variably interpretable, aspects of image quality (e.g., summary statistics of image intensities, signal-to-noise ratio, coefficient of joint variation, Euler angle, etc.) Importantly, these IQMs are defined without a *canonical* (i.e., artifact-free) reference. These IQMs can also be used for automated QC^11–14^ using machine learning models to predict scan quality ^7,15–21^.

This work proposes a protocol outlining best practices for developing and integrating QA/QC of functional, structural, and diffusion-weighted MRI (sMRI, fMRI, and dMRI, respectively) within the Standard Operating Procedures (SOPs) of whole-brain neuroimaging studies. It describes the execution of *MRIQC* to generate IQMs and visual reports designed to assess data quality. Once generated, these visual reports can be used to view and rate individual scans expeditiously. Finally, these IQMs and ratings can be used to curate datasets.

### Development of the protocol

This protocol describes how to carry out the visual assessment of T_1_-weighted (T1w), T_2_- weighted (T2w) sMRI, BOLD (blood-oxygen-level-dependent) fMRI, and dMRI scans of the human brain with the visual reports generated by *MRIQC*^7a^. We outline how to execute the software on all the images of the input dataset using typical research settings (i.e., a high-performance computing cluster). We then detail how to screen the visual reports to identify artifacts and potential quality issues. We report the usage of the *rating widget* utility, which enables rapid annotation and minimizes bookkeeping errors by allowing the ratings and annotations to be locally downloaded as JSON files. These expert annotations can additionally be submitted to the *MRIQC Web-API*^8^, a web service for crowdsourcing and sharing MRI image quality metrics and QC visual assessments. A visual abstract of the proposed protocol is given in Figure 1. Note that depending on your data collection procedures, you may run these steps in a different order, or you may run some steps multiple times.

**Figure 1.**
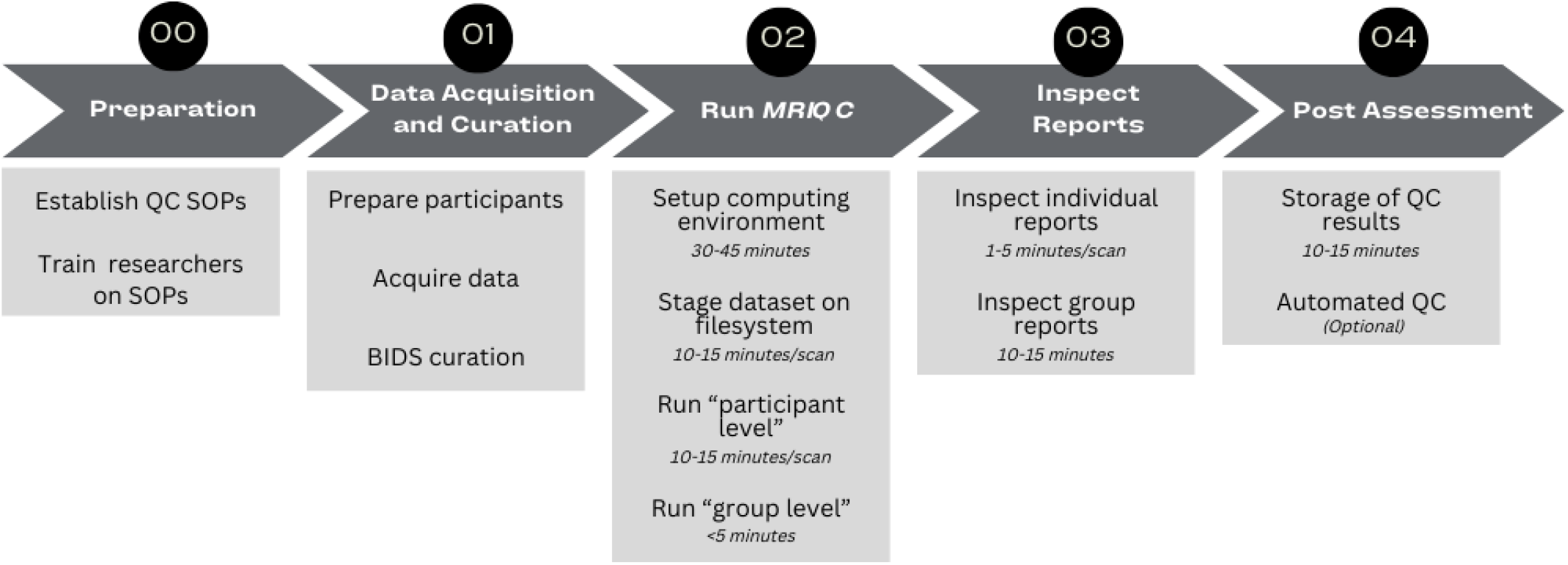
Quality Control protocol for the assessment of MRI with *MRIQC*. Step 0: Before assessing data quality, standard operating procedures (SOPs) for QC should be defined and researchers should be trained on those procedures. Step 1: After data acquisition the dataset needs to be organized following the BIDS specification. Formal verification of the BIDS structure with the *BIDS Validator* is optional but strongly recommended. Step 2: Researchers should use SLURM or other HPC job scheduler to execute *MRIQC* software on every participant for assessment. *MRIQC* generates a visual report for each of the input scans. Step 3: Researchers need to inspect those reports for quality assurance and assign a quality score in accordance with SOPs. The final rating can be shared via *MRIQC Web-API* and/or saved as a JSON text file. Step 4: Finally, users should review and store those QC reports alongside the data. Optionally, users can also train a classification or regression model on their data to operate within-site quality rating prediction to aid the assessment of future collection efforts.

As further described in the paper originally presenting the tool^22^, *MRIQC* leverages the Brain Imaging Data Structure^23^ (BIDS) to understand the input dataset’s particular features and available metadata (i.e., imaging parameters). BIDS allows *MRIQC* to configure an appropriate workflow without manual intervention automatically. To do so, *MRIQC* self-adapts to the dataset by applying a set of heuristics that account for irregularities such as missing acquisitions or runs. Adaptiveness is implemented with modularity: *MRIQC* comprises sub-workflows, which are dynamically assembled into appropriate configurations. These building blocks combine tools from widely used, open-source neuroimaging packages. The workflow engine *Nipype*^24^ is used to stage the workflows and deal with execution details (such as resource management). Then *NiReports* visual reporting system is used to create informative “reportlets”, which are atomic visualizations of the imaging data, such as mosaic views. Finally, *NiReports* composes them in the final “individual report”.

#### Applications of the protocol and target audience

A robust QA/QC strategy is vital to any neuroimaging research workflow (for instance, as shown in our tandem protocol^25^ for the case of preprocessed fMRI). However, QC of the data immediately after its acquisition and before any subsequent processing is frequently overlooked. Instead, it is merged with quality checkpoints set after data have been preprocessed or revisited only after the results have been obtained to explain failures a posteriori. While this might seem sufficient, only considering QC after analyses can enable “cherry-picking” subjects to skew analysis results. Even if QC is done earlier in an analytics workflow, before statistical analysis but after preprocessing, artifacts that render datasets unsuitable may go unnoticed. For example, brain masking can hide artifacts most evident in the background of an MRI image.

Therefore, this protocol shows how to streamline a QA/QC checkpoint immediately after image acquisition, highlighting the importance of carrying out the step before any other data inspection or computation of derived summary statistics. For example, responsive QA/QC enables decisions such as recalling a participant to repeat an otherwise failed acquisition in complex experimental designs. The protocol also proposes checklists to standardize the assessment and mechanisms to specify exclusion criteria pre-existing before the data are screened, which is critical to minimize intra- and inter-rater variabilities and to standardize QA/QC. This protocol can also be used on fully collected datasets when there may already be hundreds of subjects to screen with the use of the visual reports generated by *MIRQC*. For even larger datasets, this protocol enables data aggregation for future use in automated QC using machine learning regression and classification models.

This protocol targets MRI practitioners who routinely include sMRI, fMRI, and/or dMRI in their imaging protocols. In particular, this protocol will be of special interest to researchers starting a career in neuroimaging, such as graduate students and research coordinators who may be delegated to carry out the curation and QC of acquired data. These “newcomer” roles are likely less familiar with the impact of low data quality on subsequent analyses or what would make one particular scan “unusable” in comparison with a poor (albeit acceptable for the purpose of the study) scan. We anticipate this protocol will also be useful for PIs aiming to standardize early and reliable QC within their laboratories. This protocol can also serve as a resource to Research Directors, Engineers, and Managers of scanning centers to outsource some of the burdens in QA and early detection of scanner-related artifacts. In the long term, this protocol sets the foundations for implementing real-time QA strategies and streamlining QC within the MRI scanner pipeline. Finally, the present approach will serve for reference in the development of QA/QC protocols for other modalities, such as Positron Emission Tomography (PET).

#### Advantages and adaptations

This protocol leverages *MRIQC*, which is a widely adopted and consolidated tool as evidenced by millions of IQMs already crowdsourced by the *MRIQC Web-API* service (Fig. 4, right panel of Poldrack et al., 2024^26^), and the over 10,000 downloads of *MRIQC*’s *Docker* image. *MRIQC* is a part of the *NiPreps (****N****euro****i****maging* ***Prep****rocessing tool****s***)^*b*^ initiative, which comes along with major advantages such as a large user base, a standardized approach to development, and complementary companion QA/QC resources. *NiPreps* provide comprehensive tooling and documentation for reporting and research protocol management. For example, we have presented a previous protocol for *fMRIPrep*^25^. *NiPreps*’ *NiReports* module provides standardized visualization components that allow users to leverage knowledge and training across tools to do robust QC at multiple necessary stages. Notably, the “*SOPs-cookiecutter*” framework stands out in implementing version-controlled, collaborative, and eventually publicly shared study SOPs.

The present protocol addresses T1w, T2w, BOLD, and dMRI of the human brain, with adaptations to other imaging modalities and animal imaging existing^27^ or planned. One significant advantage is the consistency of visualizations and procedures across these modalities, decreasing the time required for researcher training and uniformizing rater experience. *MRIQC* additionally has a rating widget to rate and annotate images easily. Finally, the *MRIQC Web-API* service enables the development of machine learning models for a more reliable and semi-manually —if not fully-automated— QA/QC.

*MRIQC* has been comprehensively tested on images acquired at 1T-3T field strengths, as showcased by the millions of unique records crowdsourced by the *MRIQC Web-API* database. We are not aware of specific issues preventing *MRIQC* from performing on high-field systems. However, given the limited availability of 7T systems for human research, these users must use caution and are specifically encouraged to report issues.

##### Imaging acceleration

Echo-planar imaging (EPI) acceleration techniques such as standard in-plane acceleration or multi-band EPI sequences employed in fMRI and dMRI, are supported. Nonetheless, these acceleration techniques may introduce specific artifacts that may be difficult to assess with *MRIQC*’s reports, requiring additional QA/QC actions.

##### Animal species

*MRIQC* for rodent brains runs on anatomical and BOLD fMRI^27^. These adaptations bridge human and preclinical QA/QC protocols, and the generation of homologous IQMs may also lead to better information integration across different model species. Because rodent MRI data typically employ high-field MRI, rodent QA/QC is also key to bridging gaps towards assessing the rapidly increasing volumes of 7T MRI of the human brain. Other species, such as nonhuman primates, are not currently supported.

##### Positron Emission Tomography

Development of *MRIQC* sibling tools for other imaging modalities such as *PETQC* PET will be able to standardly replicate the framework. With the advent of simultaneous PET/MRI scanners, the upcoming integration of PET with structural and functional MRI data further emphasizes the need for consistent and multimodal QA/QC metrics and tools.

#### Limitations

##### Multi-echo BOLD fMRI

Recent versions of *MRIQC* (above 23.1.1) generate a single report for BOLD fMRI scans employing multi-echo EPI, facilitating and expediting the assessment of these images. Within each report, every echo is visualized separately. However, defining a unique set of IQMs extracted from all echos in the scan remains as an active line of development.

##### Other MRI techniques

*MRIQC* currently only runs on T1w, T2w, dMRI, and BOLD fMRI, and does not run on other modalities such as quantitative MRI data, or non-BOLD fMRI data such as ASL.

##### *MRIQC* is intended for *unprocessed* data

The purpose of *MRIQC* is to assess the quality of the original data before any processing is performed. To assess the results of processing steps, other tools must be used. We refer the reader to previously published fMRI QA/QC guidelines^10^ to see how to assess sMRI and fMRI data preprocessed by *fMRIPrep* using its visual report, and the *QSIprep* documentation^c^ *for dMRI preprocessing*.

##### *MRIQC* is not a preprocessing tool

Despite that *MRIQC*’s workflow implements standard preprocessing steps such as head-motion estimation, neither the outputs of internal processing steps nor final outcomes should be employed in downstream analysis other than to implement QC exclusion decisions.

##### Other limitations

We have evaluated the biases introduced by the process of defacing (i.e., removal of facial features from anatomical scans to protect privacy prior to data sharing) into both human ratings and automatically computed IQMs^28^. Our results indicate that it is recommended to run QA/QC on “nondefaced” data when available^29^.

#### Approaches complementing *MRIQC* in QA/QC

##### QA/QC of the Human Connectome Project (HCP)

The HCP has maintained a rigorous MRI QA/QC protocol tailored for the project^30^. For researchers following the HCP image acquisition and processing protocols, the HCP QA/QC may be a better (although mutually non-exclusive) and more comprehensive option.

##### QC protocol of the UK Biobank (UKB)

The massive scale of the UKB required an automated solution to exclude subpar images from analyses. Alfaro-Almagro et al.^18^ developed an ensemble classifier to determine image quality based on a number of *image-derived phenotypes* (e.g., 190 features for the case of T1w images). A relevant aspect of the UKB protocol is that rather than assessing/controlling for the quality of input images, the goal is to QC the outcome of the UKB preprocessing pipelines, discarding input images that will make downstream processing pipelines fail or generate results of insufficient quality.

##### Swipes for Science

Keshavan et al.^31^ proposed a creative solution to the problem of visually assessing large datasets. They were able to annotate over 80,000 two-dimensional slices extracted from 722 brain 3D images using *Braindr*, a smartphone application for crowdsourcing (https://braindr.us/). They also proposed a novel approach to the QC problem by training a convolutional neural network on *Braindr* ratings, with excellent results (area under the curve, 0.99). Their QC tool performed as well as an *MRIQC* classifier^7^ (which uses IQMs and a random forests classifier to decide which images should be excluded) on their single-site dataset. By collecting several ratings per screened entity, they were able to effectively minimize the noisy label problem with the averaging of expert ratings. Limitations of this work include the use of 2D images for annotation.

##### Healthy Brain Network Preprocessed Open Diffusion Derivatives (HBN-POD2)

Ritchie-Halford et al.^32^ also used crowdsourced QC annotations from processed diffusion images to train a deep learning classifier. They designed dmriprep-viewer^d^ and fibr^e^, two web applications that display visual reports from the preprocessed data, for experts and community scientists, respectively, to rate the images. These ratings and IQMs extracted from *QSIprep* were combined to successfully train a model predicting expert ratings from both image quality metrics and preprocessed data. In contrast to *MRIQC*, this work used preprocessed data.

##### BrainQCNet

Author MG and colleagues developed a deep learning solution to predict manual QC annotations assigned by experts^33^. For moderately-sized datasets, these tools could complement manual assessment (e.g., *MRIQC*’s visual reports) to reduce human errors and maximize inter-rater agreements. In large-scale datasets where screening of every image is not feasible, tools like *BrainQCNet* could be the only way to implement objective exclusion criteria consistent across studies, sites, and samples.

##### Fetal MRI QA/QC

Author OE and colleagues developed *FETMRQC*^34^, a tool derived from *MRIQC* and specifically tailored for fetal brain imaging. *FETMRQC* builds on *MRIQC*’s machine-learning framework to address the unique challenges of fetal imaging, mostly relating to uncontrolled fetal motion and heterogeneous acquisition protocols across clinical centers. Standardizing QA/QC in populations at risk, which typically are affected by accute data scarcity, is critical to ensure data reliability.

##### Real-time QA

Complementing *MRIQC* with online QA during scanning is strongly recommended. Indeed, real-time QA^35^ is an effective way of identifying quality issues early, shortening the time window they remain undetected, and allowing rapid reaction (e.g., repeating a particular acquisition within the session) to minimize data exclusion. In addition to visual inspection by trained technicians, there are automated alternatives such as *AFNI*’s real-time tooling^36^, or *OpenNFT*^37^. In addition to data quality monitoring, motion can be monitored during the scan using software like FIRMM^38^, so that researchers can intervene when participants have high motion.

##### Other manual QC utilities

Several alternatives for manual assessment exist, such as *MindControl*^39^, or *Qoala-T*^21^, however, they are typically designed for the assessment of surface reconstruction and other derivatives (e.g., segmentations and parcellations) extracted from T1w images. Indeed, reconstructed surfaces have been demonstrated to be a reliable proxy for some aspects of image quality of anatomical images^40^. As argued by Niso et al.^41^, researchers should include QC checkpoints at the most relevant points of the processing pipeline. In line with this recommendation, *MRIQC* should be used as an QA/QC checkpoint of unprocessed data in addition to (rather than instead of) other checkpoints at later steps (e.g., T1w preprocessing or *fMRIPrep* outputs) of the pipeline. Instead, these utilities should be used in addition to *MRIQC*.

## Materials

### Subject Data

▲**CRITICAL** The study must use data acquired after approval by the appropriate ethical review board. If the data are intended to be shared in a public repository such as OpenNeuro (■**RECOMMENDED**), the consent form submitted to the ethical review board should explicitly state that data will be publicly shared (e.g., the Open Brain consent^42^) and, if appropriate, the consent form and the data management plan must also comply with any relevant privacy laws regarding pseudo-anonymization (e.g., GDPR in the EU and HIPAA in the USA).

▲**CRITICAL** All subjects’ data must be organized according to the BIDS specification. The dataset can be validated (■**RECOMMENDED**) using the *BIDS-Validator*. Conversion to BIDS, and the BIDS-Validator steps are further described below. In this protocol, we use *ds002785* - an open dataset accessed through OpenNeuro.org^43^.

### Equipment Setup

#### MRI scanner

If the study is acquiring new data, then a standard whole-head scanner is required. *MRIQC* autonomously adapts the preprocessing workflow to the input data, affording researchers the possibility to fine-tune their MR protocols to their experimental needs and design.

#### Computing hardware

*MRIQC* can be executed on almost any platform with enough memory: conventional desktop or laptop hardware, high-performance computing (HPC), or cloud computing. Some elements of the workflow will require a minimum of 8GB RAM, although 16GB is recommended. *MRIQC* is able to optimize the workflow execution via parallelization. The use of 8-16 CPUs is recommended for optimal performance. To store interim results, *MRIQC* requires approximately 250MB per scan. For our example dataset, each subject generated approximately 2GB of interim results. This storage can be temporary, for example a”local scratch” filesystem of a compute node in HPC, which is a fast, local hard-disk that gets cleared after execution. If using other storage, these results can be removed after successfully running *MRIQC*.

#### Visualization hardware

The tools used in this protocol generate HTML reports to carry out visual quality control. These reports contain dynamic, rich visual elements to inspect the data and results from processing steps. Therefore, a high resolution, high static contrast, and widescreen monitor above 30” should be used if available (■**RECOMMENDED**). Visual reports can be opened with standard Web browsers, with *Mozilla Firefox* and *Google Chrome* being routinely tested (■**RECOMMENDED**)., Graphics acceleration support (■**RECOMMENDED**) improves report visualization.

#### Computing software

*MRIQC* can be manually installed (“bare-metal” installation as per its documentation) on *GNU/Linux, Windows Subsystem for Linux (WSL)*, and *macOS* systems, or executed via containers (e.g., using *Docker* for *Windows*). When setting up manually, all software dependencies must also be correctly installed (e.g., *AFNI*^36^, *ANTs*^44^, *FSL*^45^, *Nilearn*^46^, *Nipype*^24^, etc.). When using containers (■**RECOMMENDED**), a new container image is distributed from the *Docker Hub* service for each new release of *MRIQC*, which includes all the dependencies pinned to specific versions to ensure the reproducibility of the computational framework. Containers encapsulate all necessary software required to run a particular data processing pipeline akin to virtual machines. However, containers leverage some lightweight virtualization features of the *Linux* kernel without incurring much of the performance penalties of hardware-level virtualization. For these two reasons (reproducibility and computational performance), container execution is ■**RECOMMENDED**.

#### Report evaluation interface

*Q’kay*^47^ is a web server interface for viewing and rating *MRIQC*, and other reports generated by *NiReports* (■**RECOMMENDED)**. It collates ratings for each report in a *MongoDB* database for review. See the Q’kay documentation^f^ for installation and usage details.

## Procedure

### 0 Before starting data collection

#### 0.1 Specify rating procedure and scan exclusion criteria in the study SOPs

▲**CRITICAL** Exclusion criteria should be tailored towards a study’s specific analysis plan^10^. Since every analysis will have different requirements, there are no formal guidelines for what constitutes a “usable” scan, or a “unusable” scan, and a scan that is suboptimal for one analysis might be fine for another. However, some general qualities to assess are as follows: participant motion, presence of visually identifiable artifacts (especially artifacts that are ubiquitous across participants), and signal to noise ratio (SNR). *MRIQC*’s default “threshold” for plotting framewise displacement (FD) is set at 0.2 mm (see Figure 7), but “acceptable” levels of participant motion vary for different populations of participants. The impact of motion on data quality can be variable, often exacerbated by specific acquisition parameters and may interact with other artifacts like susceptibility distortion. While participant motion and SNR can be evaluated quantitatively, some artifacts require visual identification and may be difficult to conclusively diagnose.

Exclusion criteria should be explicitly detailed in the study SOPs^48^, which can then be referenced throughout data collection and QC, and shared to increase transparency. These SOPs can also be used to help train new researchers on the QC task. SOPs must contain QA/QC sections including, e.g., checklists of artifacts to look for, definitions for what constitutes an unusable scan, and prescriptions on how discrepancies between raters can be addressed.

SOPs can be created, managed, and shared using word processing software like *Word* or *Google Documents*. If version control is desired, GitHub^g^ can be used to host SOPs (■**RECOMMENDED**). See the *NiPreps*‘ documentation^h^ for an example of SOPs derived from the *SOPs-cookiecutter* template, with usage details.

▲**CRITICAL** If using *GitHub* or the *SOPs-cookiecutter* when creating the new repository, the researcher will very likely want to start a private repository as the study is unlikely to be publicly shareable at this point. *SOPs-cookiecutter* affords find-grained control over private information (e.g., researchers’ phone numbers or contact information for institutional resources), keeping these details separately from the SOPs and marking them as redacted placeholders if the SOPs are openly shared.

#### 0.2 Researcher training

Researchers will need to be familiar with *MRIQC*’s typical outputs and have visualized and assessed a sufficient number of *MRIQC* reports to be able to identify problematic scans and recognize common artifacts. The previously published fMRI QC guidelines^10^ and in particular its Supplementary Material^i^ can help for this training as it provides visual examples of what some problematic artifacts look like.

#### 0.3 Researcher calibration

Whenever possible, more than one researcher should assess each scan to avoid rater bias, and to ensure a robust QC process (■**RECOMMENDED**). In general, ratings and artifact categorizations do not need to be exactly the same, but large discrepancies, especially discrepancies between categorizing scans as unacceptable or acceptable, should be discussed in the context of the data analysis plans and rectified.

### 1 Data acquisition and curation

#### 1.1 Participant preparation

Data collection is an integral part of the QC process. Obtain informed consent from subjects, collect prescribed phenotypic information (sex, handedness, etc.), and prepare the participant for the scanning session. The SOPs should include a script for the interaction with the participant and a final checklist to be followed during the preparation of the experiment and setting up (■**RECOMMENDED**; see these SOPs^j^ for an example). Due to the negative impact of motion on data, participants should be thoroughly informed of the importance of staying still, and care should be taken to ensure padding is placed properly for their comfort. Additionally, for populations where increased motion or scan discontinuation due to discomfort or anxiety is likely (children, elderly, some clinical populations), a mock scanner should be utilized prior to the acquisition scan, so that the participants can get acclimated to the scanner bore. Participants should also be informed of the optimal time for swallowing and adjusting position prior to any scans.

#### 1.2 MRI acquisition

Run the prescribed scan protocol. Between acquisitions, continue to check in on participants and remind them to stay still. Researchers may want to visually monitor motion during the scan, especially for high motion populations. Use *ReproIn*^49^ naming conventions when defining the prescribe sequences to ease later conversion to BIDS (■**RECOMMENDED**).

▲**CRITICAL** Keep a pristine copy of the original data and metadata.

#### 1.3 BIDS conversion

Store all imaging data in NIfTI-1 or NIfTI-2 file formats as per BIDS specifications, ensuring all metadata are correctly encoded. The process can be made much more reliable and consistent with conversion tools such as *dcm2niix*^50^ or *HeuDiConv*^45^. The *ReproIn*^18^ naming convention automates the conversion to BIDS with *HeudiConv*, ensuring the shareability and version control of the data starting from the earliest steps of the pipeline. For larger datasets with heterogeneous acquisition parameters *CuBIDS (“Curation of BIDS”)*^52^ can be used to identify all permutations of acquisition parameters.

▲**CRITICAL** If data are to be shared publicly, they must be anonymized^53^ and facial features must be removed from the anatomical images (some tools and recommendations are found with the Open Brain consent project^42^ and White et al.^54^). Despite defacing, it is ■ **RECOMMENDED** to execute *MRIQC* on the original images before defacing (“nondefaced”). In such cases, maintaining a “private” copy of the dataset in BIDS will be necessary.

To ensure that the dataset is BIDS-compliant, use the online BIDS-Validator^k^ or some up-to-date local native or containerized installation^l^, specifying the path to the top-level directory of the dataset (■**RECOMMENDED**). The online BIDS-Validator can be run in any modern browser without uploading any data.

### 2 Execute *MRIQC*

The protocol is described assuming that execution takes place on an HPC cluster with the *Bash* shell, the *SLURM* job scheduler^55^ and the *Apptainer* container framework^56^ (v3.0 or higher) installed. With appropriate modifications to the batch-submission directives, the protocol can also be deployed on HPC clusters with alternative job management systems such as *SGE, PBS* or *LSF*. For execution in the cloud or on conventional desktop or laptop, please refer to *MRIQC’s* documentation^m^.

#### 2.1 Setting up the computing environment (⬤TIMING 30-45 min)

First, define `$STUDY`, an environment variable pointing at the directory containing all study materials:

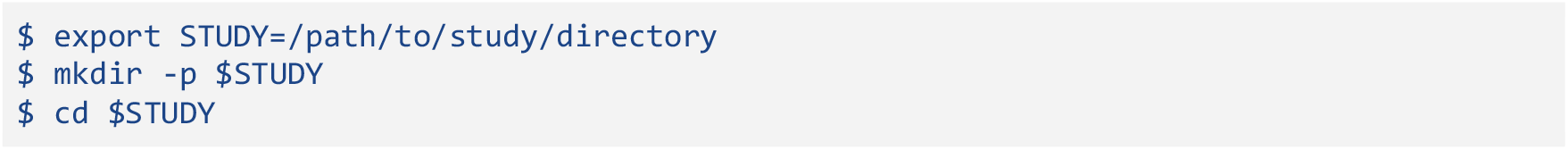

When running *MRIQC* for the first time in a new computing environment, begin by creating a container image. If this is being done on an HPC, be sure to use a compute node rather than a login node to avoid your process being killed for using too many resources. As of *Apptainer* 2.5, it is straightforward to download the container image via the Docker registry:

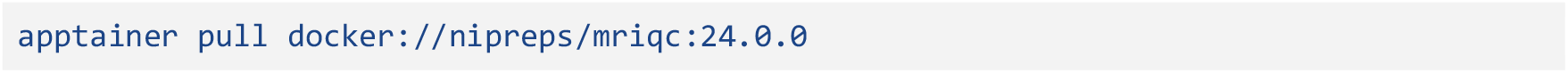

This command line will create an Apptainer image file at the current working directory (`$STUDY/mriqc_24.0.0.sif`).

**▲CRITICAL** Be sure to indicate a specific version of *MRIQC* (version 24.0.0, in this example, but the most up to date *MRIQC* is ■**RECOMMENDED**). The quality of all datasets and subjects used in any study should be assessed consistently, using the same version of *MRIQC*. The version of *MRIQC* previously used to process any dataset can be identified by consulting the `GeneratedBy` field of the `dataset_description.json` file in the top level of *MRIQC*’s output directory or by consulting the `MRIQC version` field in the `Summary` box at the top of any visual reports.

#### 2.2 Stage the dataset on the computing platform (⬤TIMING 15 min)

Transfer a copy of the nondefaced (if available) BIDS dataset to the designated filesystem that will be accessible from compute nodes. In this protocol case, this path will be represented by the environment variable `$STUDY`.

If you’re using data that you’ve collected, you can copy, move, or download your dataset into `$STUDY`. If you’re using data available from *OpenNeuro* you can download it using the code snippets provided on *OpenNeuro* for Amazon Web Service S3 or *Datalad*.

■ **RECOMMENDED** To download the example dataset used in this protocol, deploy a lightweight *Python* environment in the cluster using *Conda, Anaconda, Miniconda*, or *Mamba* and install *DataLad* and its dependency *git-annex*.

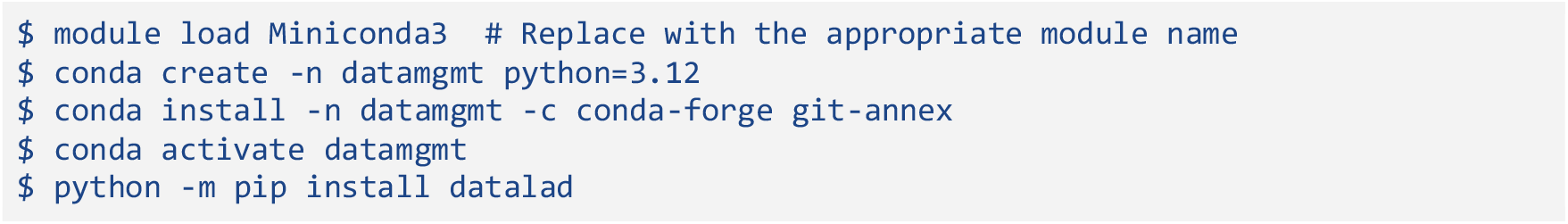

*DataLad* can be used for more advanced dataset management, but for this particular protocol, *DataLad* is only used to download a dataset from *OpenNeuro*.

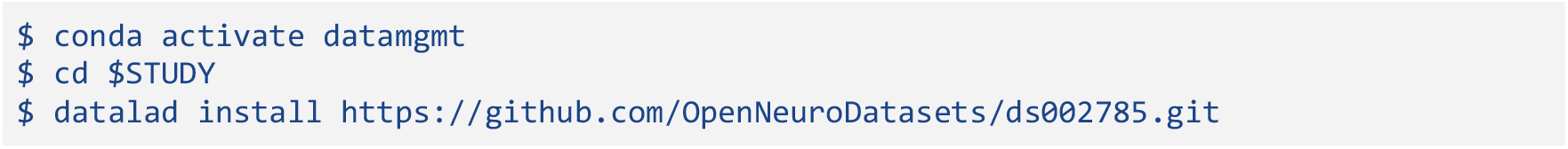

When downloading *DataLad* datasets as above, *DataLad* will not download large imaging files onto the hard-disk, only the dataset structure and human-readable textual files. *MRIQC* integrates *DataLad* starting with its 22.0.3 release and will transparently pull images from the remote storage locations as they are submitted for processing, minimizing storage needs for large datasets. In order to use *DataLad* to manage other datasets from scratch, refer to the *DataLad* Handbook^n^ for further information and reference.

#### 2.3 Run *MRIQC*’s “participant” level (⬤TIMING 10-15 minutes per scan, depending on the number, length, and size of imaging schemes in the protocol)

Container instances can make use of multiple CPUs to accelerate subject level processing, and multiple container instances can be distributed across compute nodes to parallelize processing across subjects (■**RECOMMENDED**). To run *MRIQC*, first create a batch prescription file with the preferred text file editor, such as *Nano* (`nano $STUDY/mriqc.sbatch`). Box 2 describes an example of a batch prescription file `$STUDY/mriqc.sbatch`, and the elements that may be customized for the particular execution environment. After adapting the example script to your HPC using your preferred text editor, submit the job to the scheduler: `sbatch $STUDY/mriqc.sbatch`. Although the default options are typically sufficient, the documentation of *MRIQC*^o^ provides more specific guidelines.

#### 2.4 Run *MRIQC*’s “group” level (⬤TIMING >5 min)

Once all “participant” level jobs have completed, run the “group” level using the same paths and commands defined in Box 2 to aggregate IQMs and generate the group-level report.

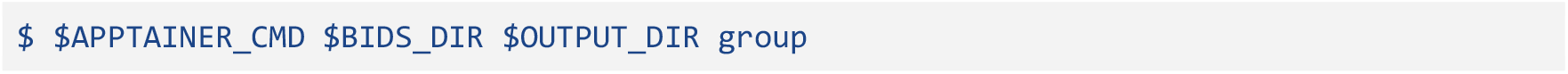

### 3 Visual inspection of reports

#### 3.1 Inspect all visual reports generated by *MRIQC* (⬤TIMING 1-5 min per scan)

After running *MRIQC*, inspect all the generated visual reports to identify images with insufficient quality for analysis, according to predefined exclusion criteria. Each scan gets an HTML report, consisting of “reportlet” visualizations that highlight a specific aspect of the scan quality. Refer to the shared reports^p^ for examples, Box 3, Box 4, or Box 5 for an inspection protocol, and refer to up-to-date [documentation](https://mriqc.readthedocs.io/en/latest/reports.html). For interested readers, more details about the artifacts to inspect, notably the explanation behind their emergence and how to differentiate artifacts that look similar, can be found in the fMRI QC guidelines^10^. The built-in rating widget should be used to record overall image quality rating and specific artifacts (Figure 2). *Q’kay* can be used to manage reports and ratings (■**RECOMMENDED**). To minimize bias, reports should be viewed in a random order, and environmental variables (such as screen brightness or size) should be consistent.

**Figure 2.**
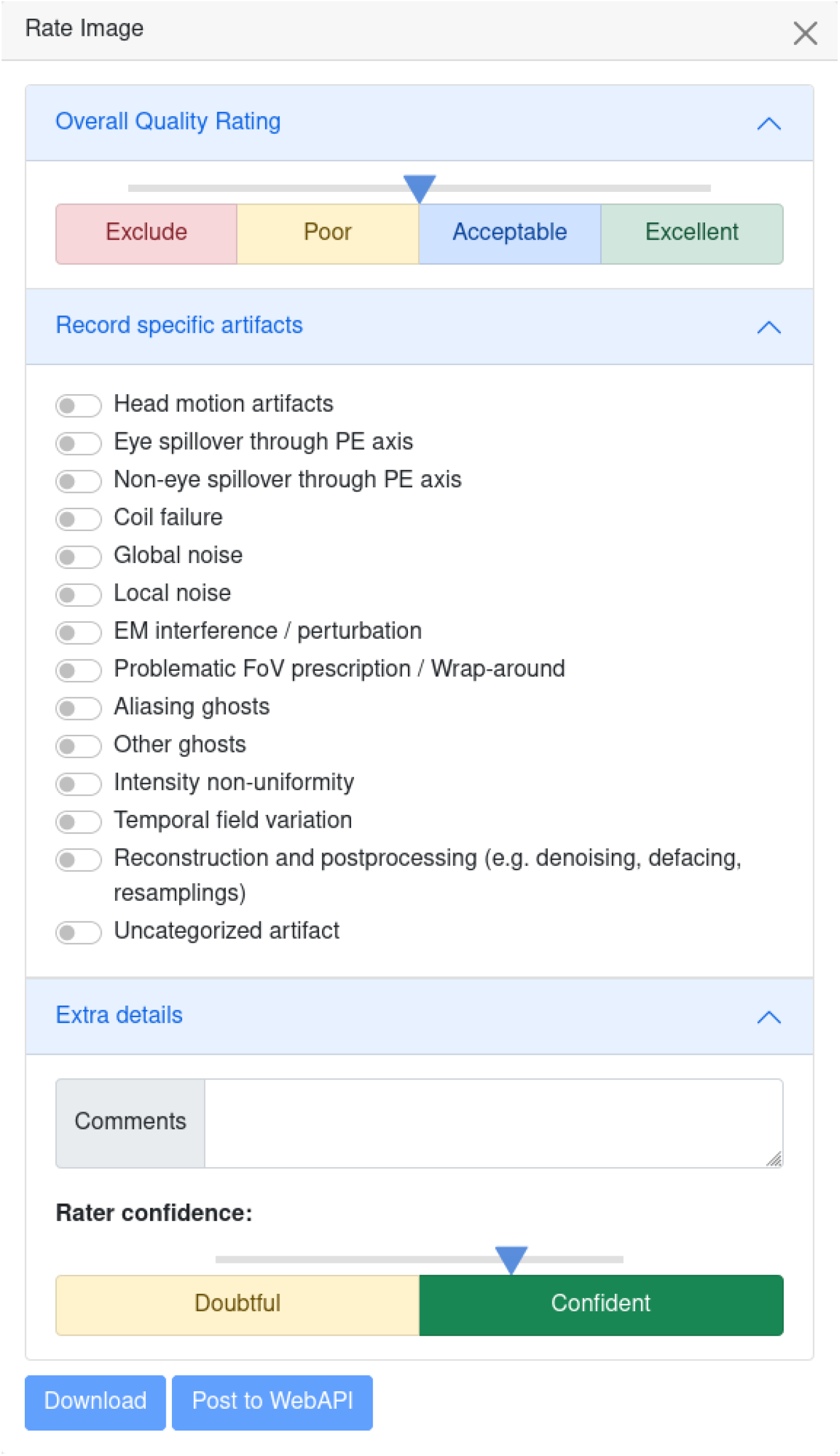
Ratings Widget With the ratings widget, you can take note of which artifacts are present in the scan, and give it an overall quality score using the slider ranging on a continuous scale from 1 to 4 (1 : exclude, 4 : excellent quality). This information will be locally downloaded as a JSON file, which can be saved along with other dataset metadata, and used for final dataset curation. This data can also optionally be uploaded to the MRIQC Web-API for crowdsourced QC projects.

Some examples of artifacts that could grant exclusion of images from a study are T1w images showing extreme ringing as a result of head motion, irrecoverable signal dropout derived from susceptibility distortions across regions of interest in fMRI or dMRI, excessive N/2 ghosting within fMRI scans, or excessive signal leakage through slices in multiband fMRI reconstructions. Some artifacts may be more obvious in certain visualizations and more subtle in others, so inspecting reports is not strictly a linear process (i.e., once an artifact is identified in one component, checking other components for evidence can help diagnose the problem or evaluate the severity).

Visualizations generated with the `--verbose-reports` flag should be used for debugging software errors, rather than for evaluating scan quality. *MRIQC* implements a quick and coarse workflow, so those visualizations checking the validity of intermediate steps should not be considered to evaluate overall quality.

##### Box 2.

###### Running *MRIQC* on HPC

Execution of BIDS-Apps^57^ (such as *MRIQC* or *fMRIPrep*^58^) is easy to configure on HPC clusters. We provide below an example execution script for a SLURM-based cluster. An up-to-date, complete version of the script is distributed within the documentation^q^.

~~~
#!/bin/bash
##NOTE: These should work with Slurm HPC systems,
 # but these specific parameters have only been tested on
 # Stanford’s Sherlock. Some parameters may need to be
 # adjusted for other HPCs, specifically --partition.
#SBATCH --job-name mriqc
#SBATCH --partition normal #TODO: update for your HPC
#NOTE: The --array parameter allows multiple jobs to be launched at once,
 # and is generally recommended to efficiently run several hundred jobs
 # at once.
##TODO: adjust the range for your dataset; 1-n%j where n is the number of
 # participants and j is the maximum number of concurrent jobs you’d like
 # to run.
#SBATCH --array=1-216%50
#SBATCH --time=1:00:00 #NOTE: likely longer than generally needed
#SBATCH --ntasks 1
#SBATCH --cpus-per-task=16
#SBATCH --mem-per-cpu=4G
# Outputs ----------------------------------
#SBATCH --output log/%x-%A-%a.out
#SBATCH --error log/%x-%A-%a.err
#SBATCH --mail-user=%u@stanford.edu #TODO: update for your email domain
#SBATCH --mail-type=ALL
# ------------------------------------------
STUDY=“/scratch/users/mphagen/mriqc-protocol” #TODO: replace with your path
MRIQC_VERSION=“24.0.2” #TODO: update if using a different version
BIDS_DIR=“${STUDY}/ds002785” # TODO: replace with path to your dataset
OUTPUT_DIR=“${BIDS_DIR}/derivatives/mriqc-${MRIQC_VERSION}”
APPTAINER_CMD=“apptainer run -e mriqc_${MRIQC_VERSION}.sif”
# Offset subject index by 1 because of header in participants.tsv
subject_idx=$((${SLURM_ARRAY_TASK_ID} + 1))
##NOTE: The first clause in this line selects a row in participants.tsv
 # using the system generated array index variable SLURM_ARRAY_TASK_ID.
 # This is piped to grep to isolate the subject id. The regex should
 # work for most subject naming conventions, but may need to be modified.
subject=$(sed -n ${subject_idx}p ${BIDS_DIR}/participants.tsv \
 | grep -oP “sub-[A-Za-z0-9_]*”)
echo Subject $subject
cmd=“${APPTAINER_CMD} ${BIDS_DIR} ${OUTPUT_DIR} participant \
  --participant-label $subject \
  -w $PWD/work/ \
  --omp-nthreads 10 --nprocs 12”
echo Running task ${SLURM_ARRAY_TASK_ID}
echo Commandline: $cmd
eval $cmd
exitcode=$?
echo “sub-$subject ${SLURM_ARRAY_TASK_ID} $exitcode” \
>> ${SLURM_ARRAY_JOB_ID}.tsv
echo Finished tasks ${SLURM_ARRAY_TASK_ID} with exit code $exitcode
exit $exitcode
~~~

##### Box 3.

###### Basic visual report

The visual report consists of various detailed visualizations of the raw data. We detail below which reportlets are presented for anatomical images and explain potential pitfalls in each visualization.

**Figure 3.**
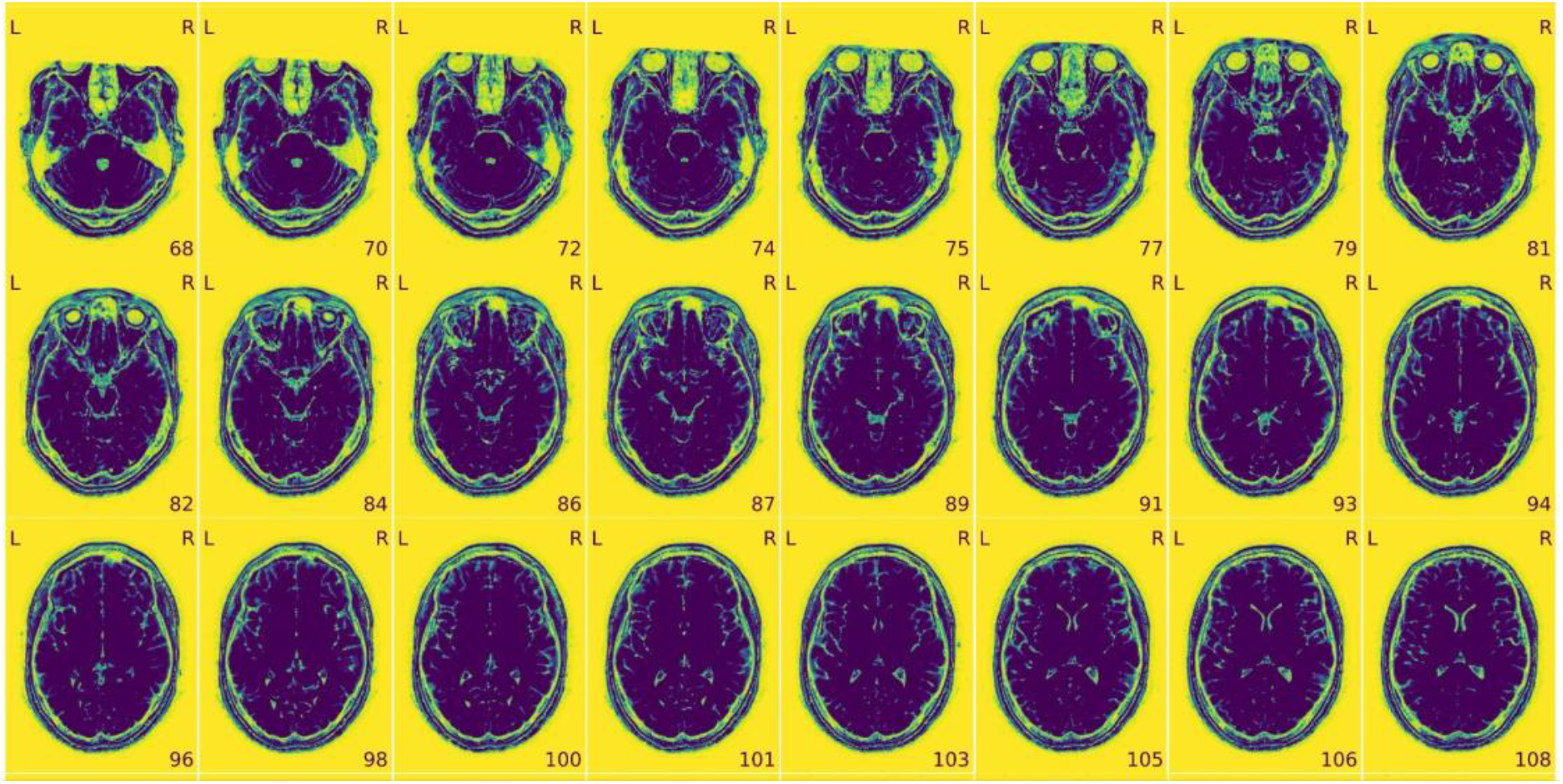
View of the background of the anatomical image. This visualization is an enhancement of the background noise. Any artifacts identified in this image may also be visible in the zoomed-in image. Look for: • Excessive visual noise in the background of the image, specifically in or around the brain, or “structured” noise (global or local noise, image reconstruction errors, EM interference). • Motion artifacts, commonly seen as “waves” coming out from the skull (head motion artifacts). • Faint and shifted copies of the brain in the background (aliasing ghosts).

**Figure 4.**
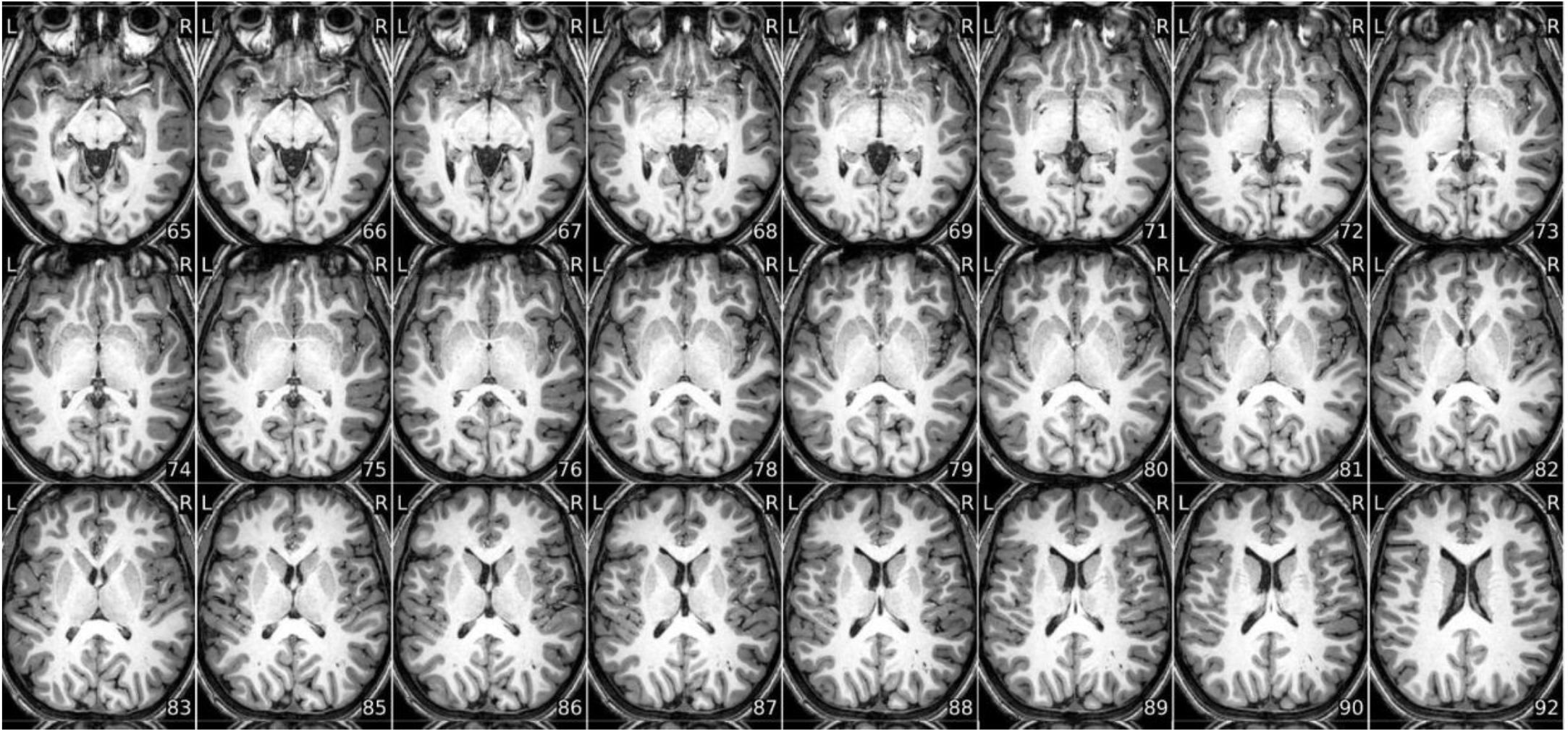
Zoomed-in mosaic view of the brain. This visualization is the T1w or T2w scan, sliced, and plotted in a mosaic view, allowing you to look at the whole brain at once. Look for: • Brain not displayed in the right orientation (axis flipped or switched because of data formatting issues). • Ripples caused by head motion or Gibbs ringing that blurs some brain areas. • Brain areas that are distorted, or extreme deviations from typical anatomy, which may indicate an incidental finding, or an artifact such as susceptibility distortion. • Inconsistency in the contrast between light and dark areas of the brain that does not reflect actual gray matter/white matter anatomy. It is typically caused by intensity non-uniformity, which manifests as a slow and smooth drift in image intensity throughout the brain. • “Wrap-around” where a piece of the head gets cut-off and folded over on the opposite extreme of the image. It is a problem only if the folded region contains or overlaps with the region of interest (problematic FOV). • Eye-spillover, where eye movements trigger signal leakage that might overlap with signal from regions of interest (eye spillover through PE axis).

###### About

Contains metadata (filename, report creation date and time), *MRIQC* version information and specific workflow details. May contain Warnings or Errors, which can be searched on NeuroStars.org^r^ or the *MRIQC* Github Repository^s^.

###### Extracted Image Quality Metrics (IQMs)

Metrics calculated by *MRIQC* that relate to the quality of the images. These are also available in each subject’s directory as a JSON file. Definitions for each IQM can be found in the *MRIQC* documentation^t^.

###### Metadata

Scan metadata from the BIDS sidecar such as RepetitionTime, EchoTime and ScannerSequence.

###### Provenance information

Information related to the version of *MRIQC* used to generate the visual report.

##### Box 4.

###### Basic echo-wise reports

The report once again consists of various detailed visualizations of the raw data. For each reportlet presented, we detail below quality aspects for careful consideration. If the data contains multiple echoes, each echo is visualized separately for these reportlets.

**Figure 4.**
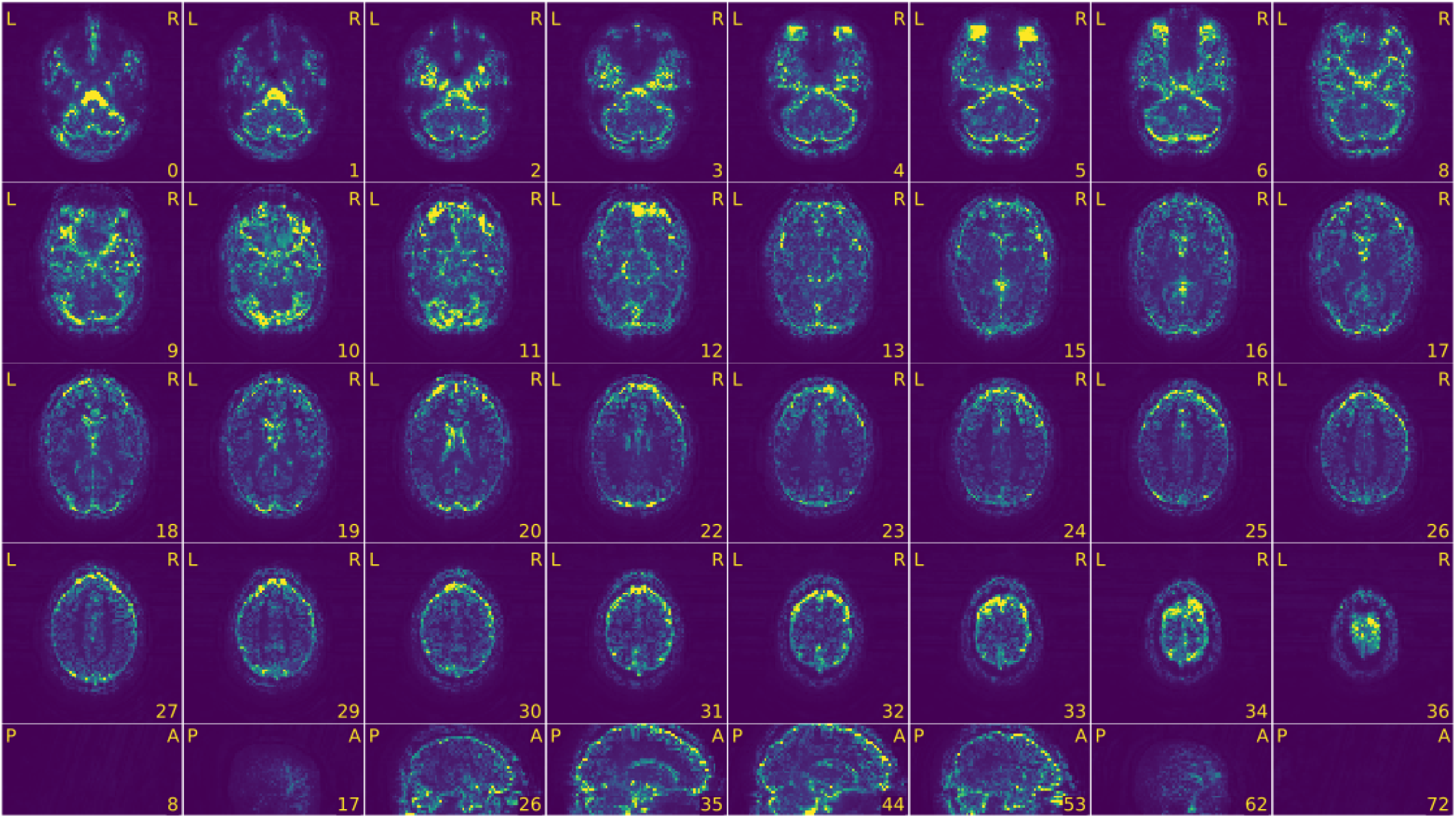
Standard deviation of signal through time. This visualization corresponds to the standard deviation of the BOLD signal in each voxel, plotted as sagittal and axial cross sections, with yellow representing the most extreme values. The eyes and arteries will typically be the brightest yellow, a result of physiological motion. Look for: • An extra outline of the brain shifted on the acquisition-axis (aliasing ghost if it overlaps with the actual image, other ghost if it does not). • Piece of the head (usually the front or back) outside of the field of view that folds over on the opposite extreme of the image (problematic FOV prescription / wrap around). This is a problem only if the folded region contains or overlaps with the region of interest. • Symmetric brightness on the edges of the skull (head motion). • Any excessive variability as indicated by patterns of brightness that is unlikely to be genuine BOLD activity, such as vertical strikes in the sagittal plane that extends hyperintensities through the whole plane.

The severity of artifacts identified in the standard deviation map can be evaluated by looking for evidence of them in the BOLD average or raw NifTI.

**Figure 5.**
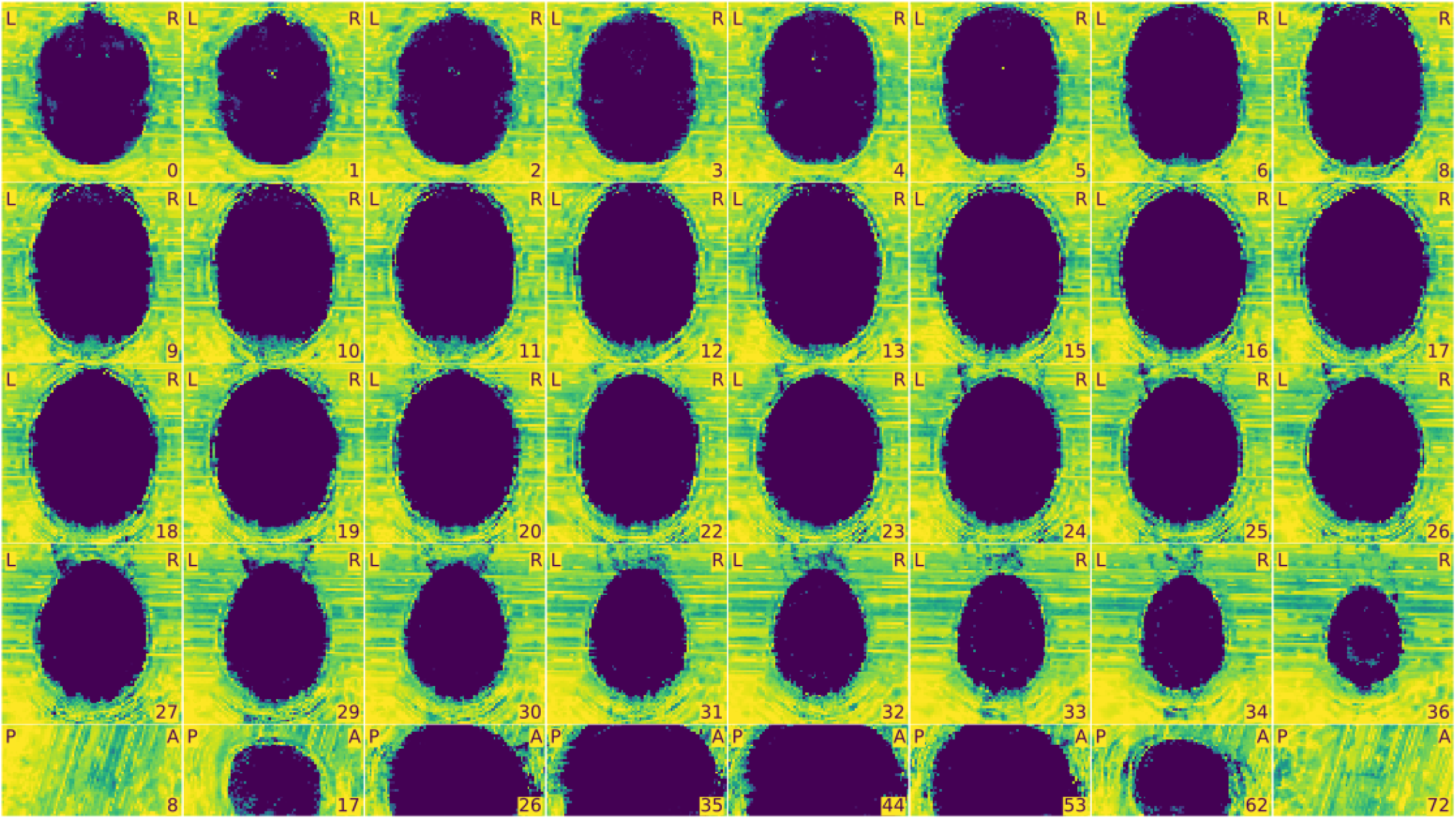
View of the background of the voxel-wise average of the BOLD timeseries. Mosaic view of the average BOLD signal, with background enhancement. Look for: • An extra outline of the brain shifted on the acquisition-axis (aliasing ghost if it overlaps with the actual image, other ghost if it does not). • Excessive visual noise in the background of the image, specifically in or around the brain, or “structured” noise (global or local noise, image reconstruction errors, EM interference).

**Figure 6.**
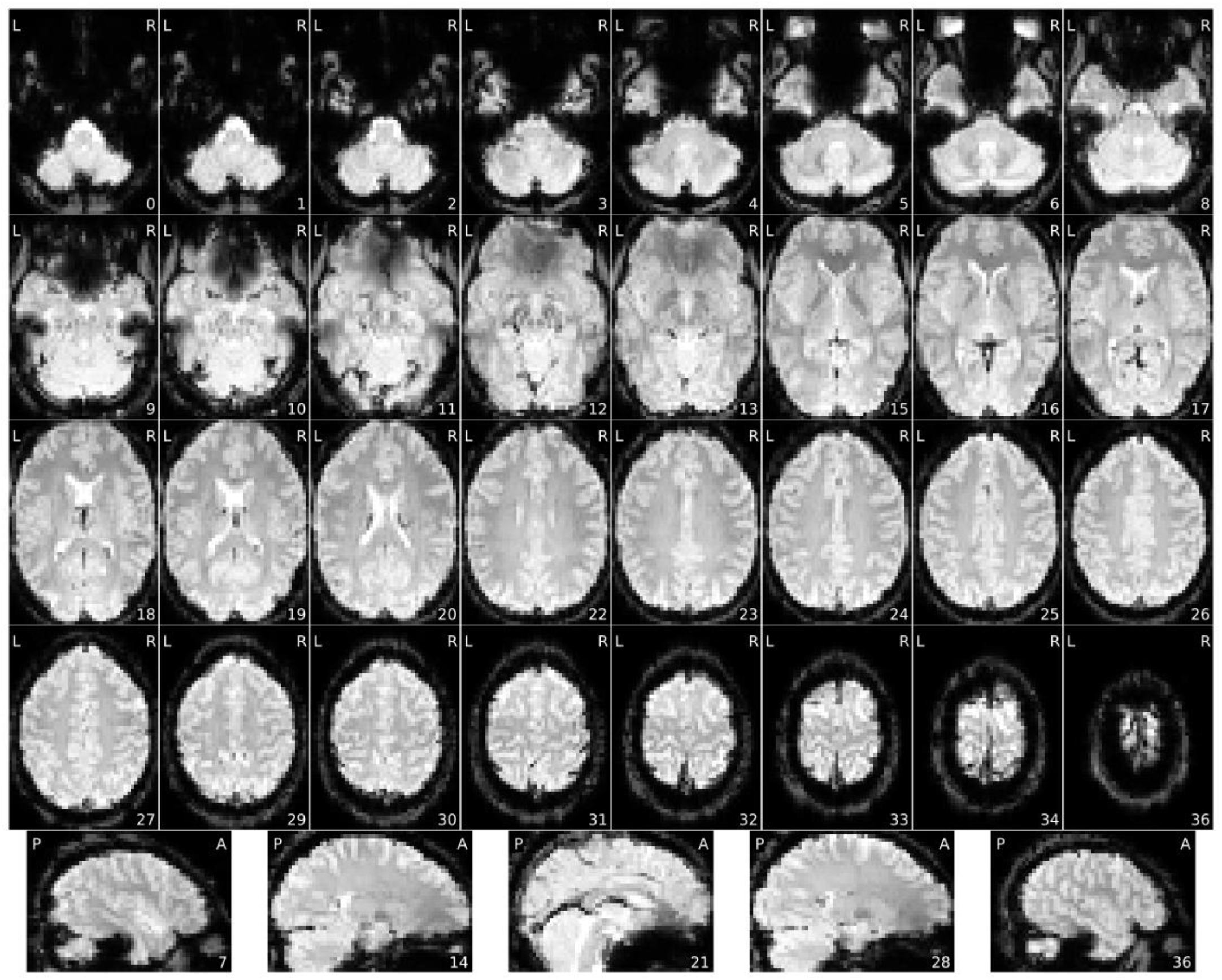
Voxel-wise average of BOLD time-series, zoomed-in covering just the brain. This visualization is the average values of the BOLD signal in each voxel across the entire scan duration, plotted as sagittal and axial cross sections. Look for: • Brain not displayed in the right orientation (axis flipped or switched because of data formatting issues). • Piece of the head (usually the front or back) outside of the field of view that folds over on the opposite extreme of the image (problematic FOV prescription / wrap around). It is a problem only if the folded region contains or overlaps with the region of interest. • Missing or particularly blurry slices (coil failure, local noise). • Signal drop-outs or brain distortions, especially close to brain/air interfaces such as prefrontal cortex or the temporal lobe next to the ears (susceptibility distortions), or from unremoved metallic items (EM interference). • Uneven image brightness, especially near anatomical areas that would be closer to the head coils (intensity non-uniformity). • Blurriness (local noise if confined to one area of the scan, global noise if present in the entire image).

**Figure 7.**
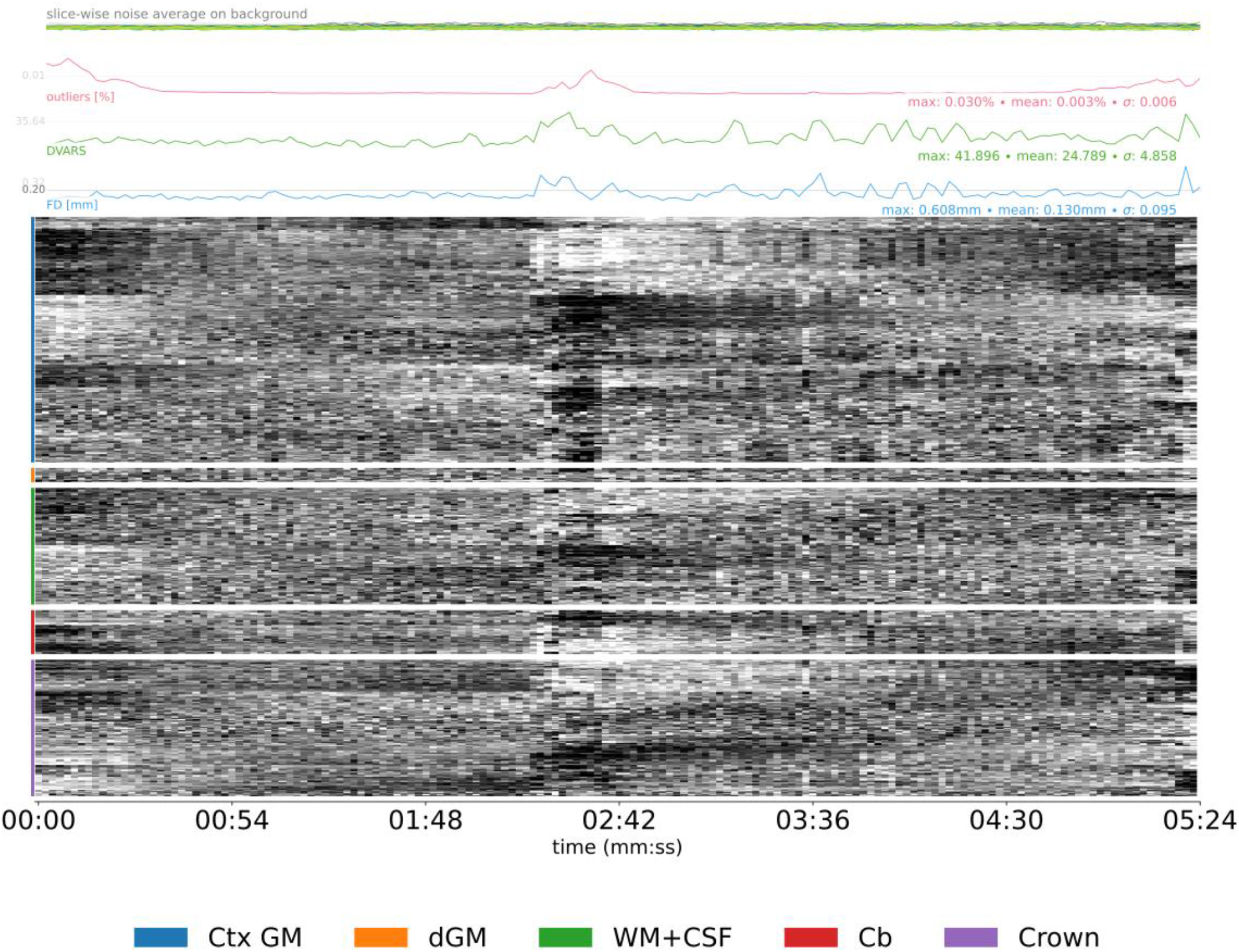
Carpetplot and nuisance signals. This reportlet is aimed at emphasizing changes in voxel intensity throughout an fMRI scan. The core of this visualization is the carpet plot, which works by plotting voxel time series in close spatial proximity so that the eye notes temporal coincidence^59^. The carpet plot is segmented into relevant regions, notably the “crown” or brain-edge area that corresponds to voxels located on a closed band around the brain^60^. Above the carpet plot, several time series are represented to support the interpretation of the carpet. The time series plotted are the slice-wise noise average on background, the % outliers, DVARS, and framewise displacement (FD). Look for: • Prolonged dark deflections on the carpet plot paired with peaks in FD (motion artifacts caused by sudden movements). • Periodic modulations of the carpet plot (motion artifacts caused by regular, slow movement like respiration). • Sudden change in overall signal intensity on the carpet plot not paired with FD peaks, and generally sustained through the end of the scan (coil failure). • Strongly polarized structure in the crown region of the carpet plot is a sign of artifacts, because those voxels should not present signal as they are outside the brain. • Consistently high values and large peaks in FD or DVARS (motion artifacts). • An average FD above your pre-defined threshold (motion artifacts). • Sudden spikes in the slice-wise noise average on background that affect a single slice (motion artifact or white-pixel noise depending on whether it is paired with a peak in FD or DVARS or not).

###### About

Identical to Box 4 “About”.

##### Box 5.

###### Summary

**Figure 8.**
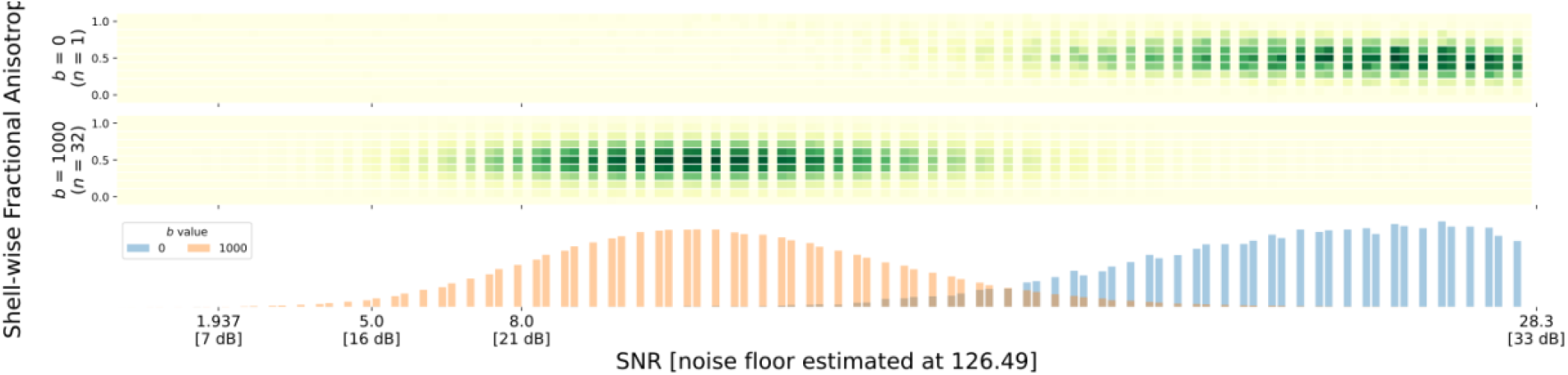
Shell-wise joint distribution of SNR vs FA in every voxel. The top visualization shows a heatmap of the estimated SNR and FA for every voxel by shell. The higher shells should have lower SNRs. The bottom visualization shows the histogram of SNR values independent of FA, separated by shell. Look for: • Overlap between the SNR distributions for each shell (can indicate suboptimal acquisition parameters). • Linear correlations between FA and SNR, which indicates noise contamination. • Non-normal distributions of SNR for each shell; MRI noise is Rician, which means that SNR distributions that are mostly above approximately 2 should resemble a normal distribution in each shell^61^.

**Figure 9.**
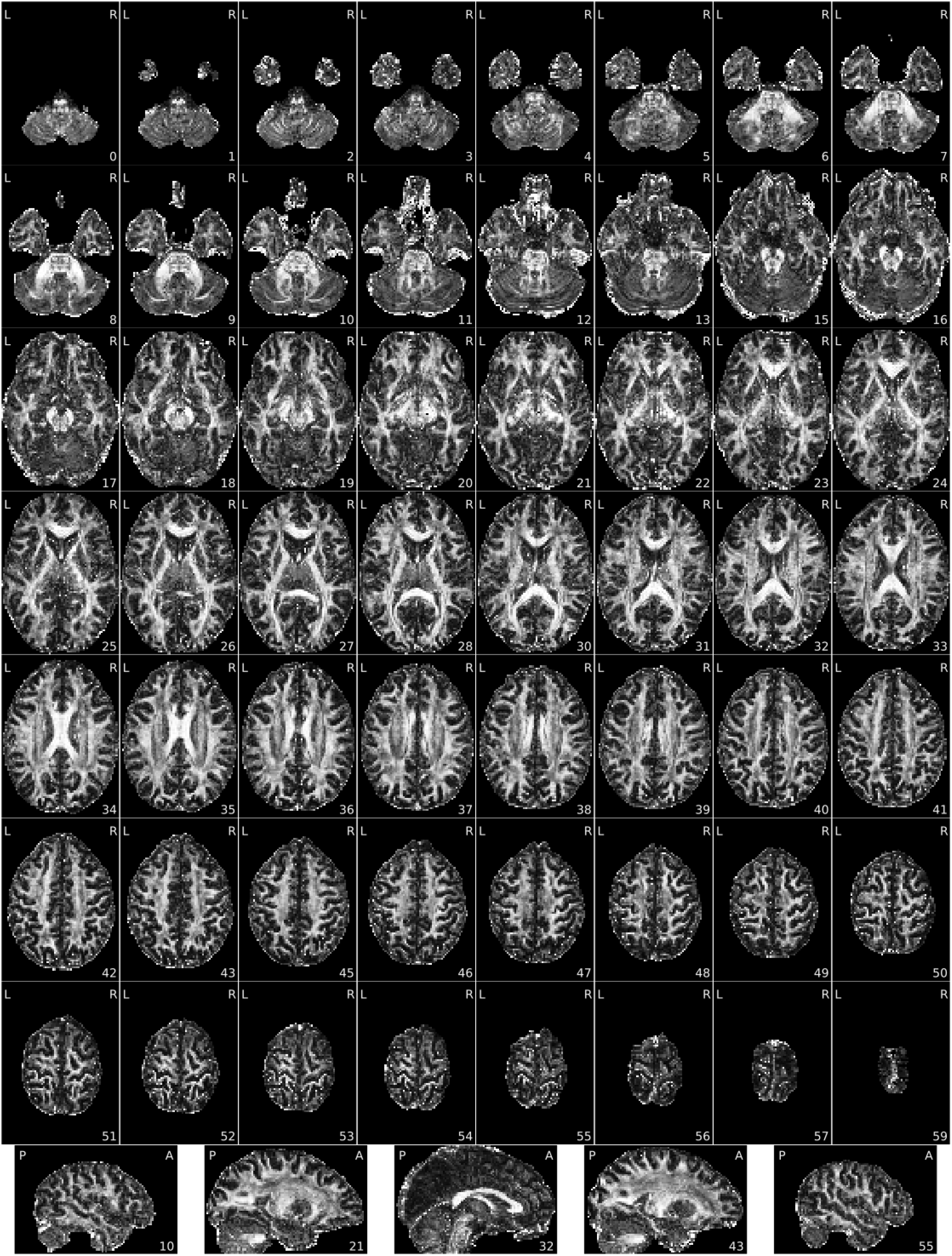
Fractional anisotropy (FA) map. The reconstructed FA values for each voxel, plotted by slices in a mosaic layout. Look for: • Brain not displayed in the right orientation (axis flipped or switched because of data formatting issues). • Piece of the head (usually the front or back) outside of the field of view that folds over on the opposite extreme of the image (problematic FOV prescription / wrap around). It is a problem only if the folded part overlaps with the region of interest. • Blurriness or lack of contrast between white matter and gray matter. You should be able to identify major white matter structures, such as the corpus callosum. FA should be higher in white matter tissue, and especially high in white matter areas where there is a predominant fiber orientation (e.g., the corpus callosum)., • Excessive white speckles, especially if they’re located in the interior of the brain (reconstruction error).

**Figure 10.**
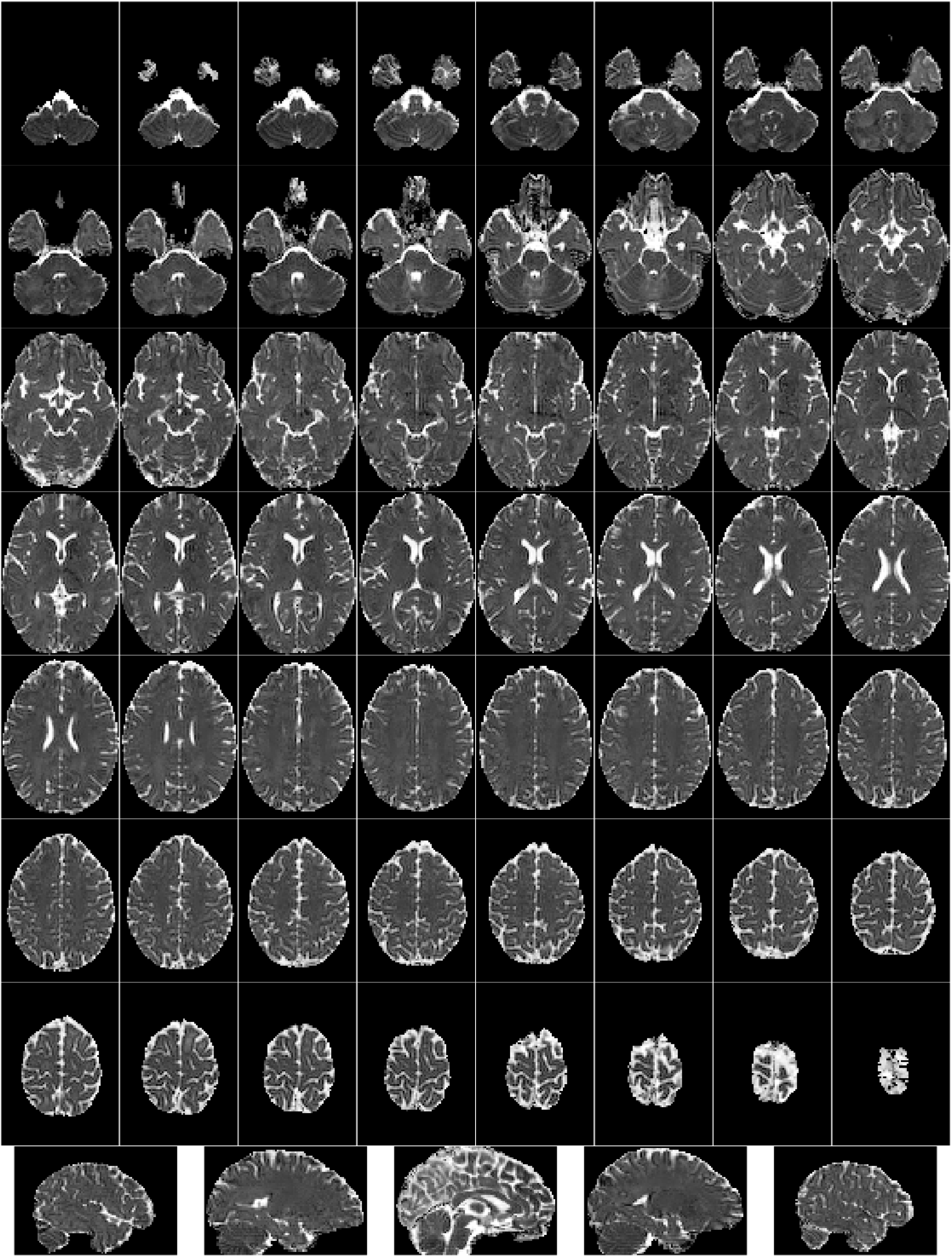
Mean diffusivity (MD) map. The reconstructed MD values for each voxel, plotted by slices in a mosaic layout. Look for: • Brain not displayed in the right orientation (axis flipped or switched because of data formatting issues). • Piece of the head (usually the front or back) outside of the field of view that folds over on the opposite extreme of the image (problematic FOV prescription / wrap around). • Blurriness or lack of contrast between white matter, gray matter, and ventricles (local noise if confined to one area of the scan, global noise if present in the entire image).

###### DWI shells

The following reportlets are created for each shell.

**Figure 11.**
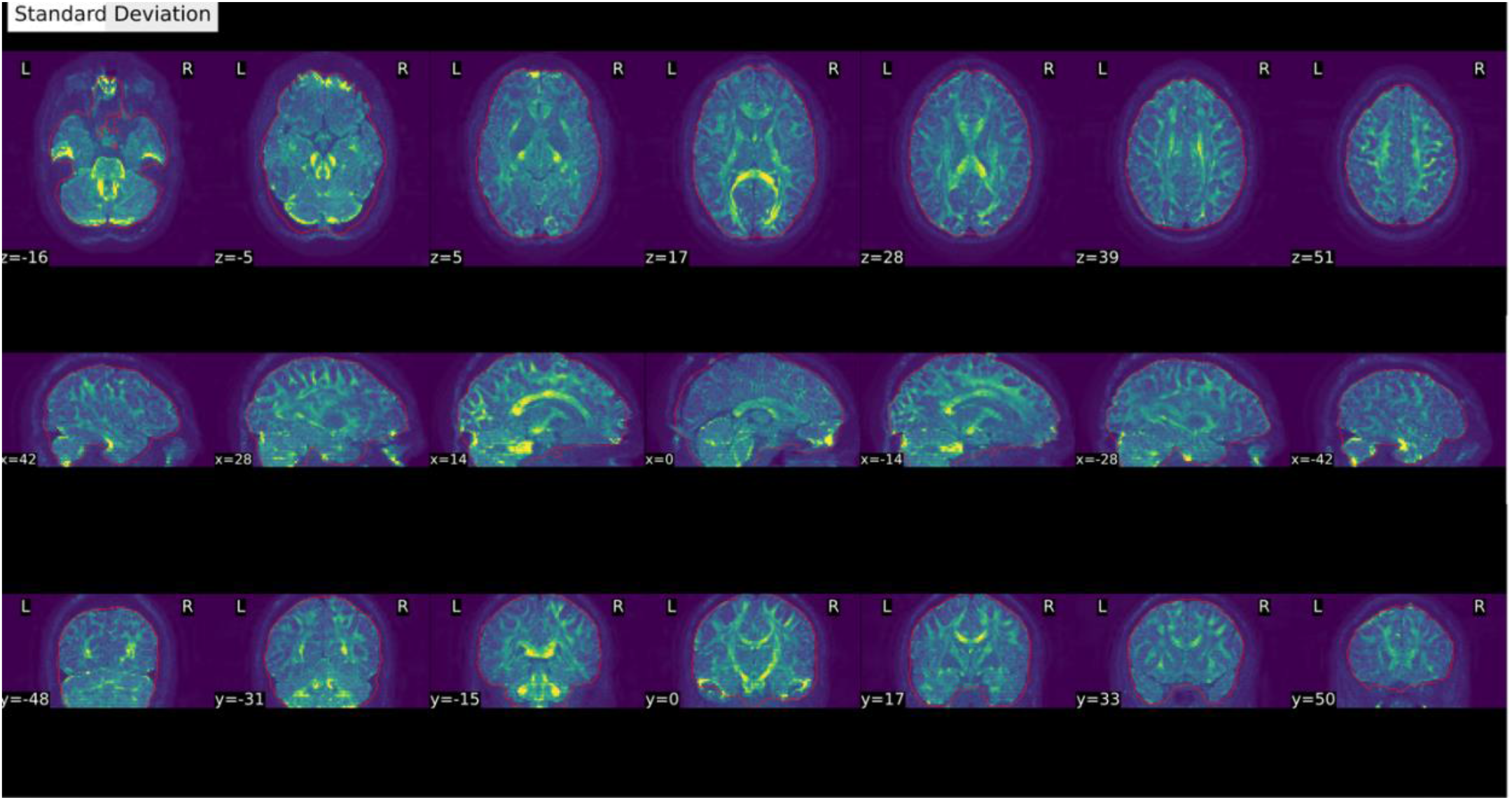
Voxel-wise average and standard deviation across volumes in this DWI shell. This visualization transitions between the voxel-wise average and the standard deviation values for each voxel, with yellow indicating higher variance. A red line delineates the calculated brain mask. As the b-value increases, the yellow signal should clearly resemble white matter tracts. Note that shells with a b-value of 0 may not have any variability if only one gradient was collected. Look for: • An extra outline of the brain shifted on the acquisition-axis (aliasing ghost if it overlaps with the actual image, other ghost if there’s no overlap). • Piece of the head (usually the front or back) outside of the field of view that folds over on the opposite extreme of the image (problematic FOV prescription / wrap around). This is a problem only if the folded region contains or overlaps with the region of interest. • Symmetric brightness on the edges of the skull (head motion). • Any excessive variability as indicated by patterns of brightness, such as brightness localized to one section of the brain or outside of white matter tracts (global or local noise).

**Figure 12.**
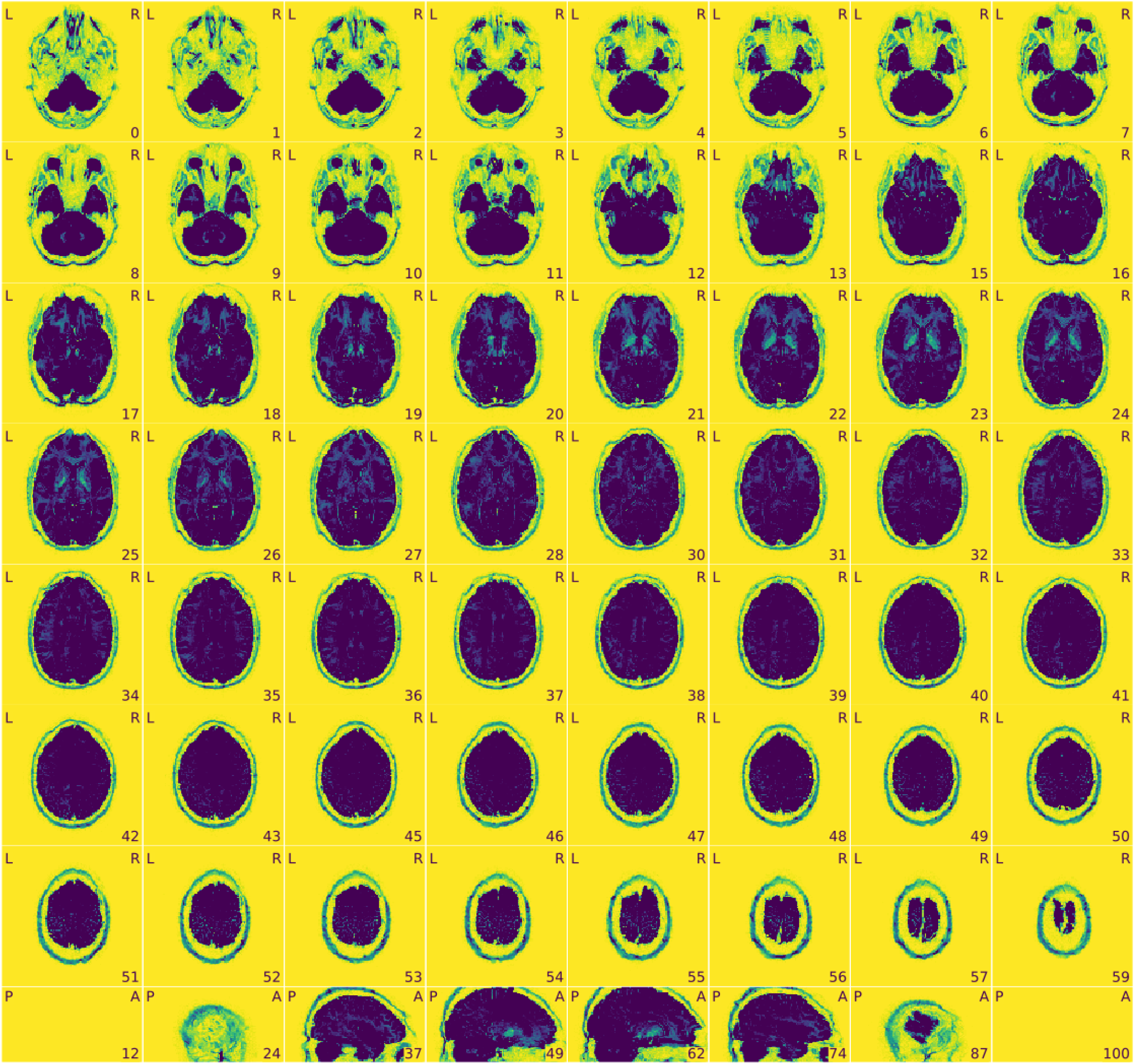
View of the background of the voxel-wise average of this DWI shell. This visualization is an enhancement of the background noise. Any artifacts identified in this image may also be visible in the FA or MD maps. Look for: • An extra outline of the brain shifted on the acquisition-axis (aliasing ghost if it overlaps with the actual image, other ghost if it does not). • Excessive visual noise in the background of the image, specifically in or around the brain, or “structured” noise (global or local noise, image reconstruction errors, EM interference).

###### About

Identical to Box 4 “About”.

**Figure 13.**
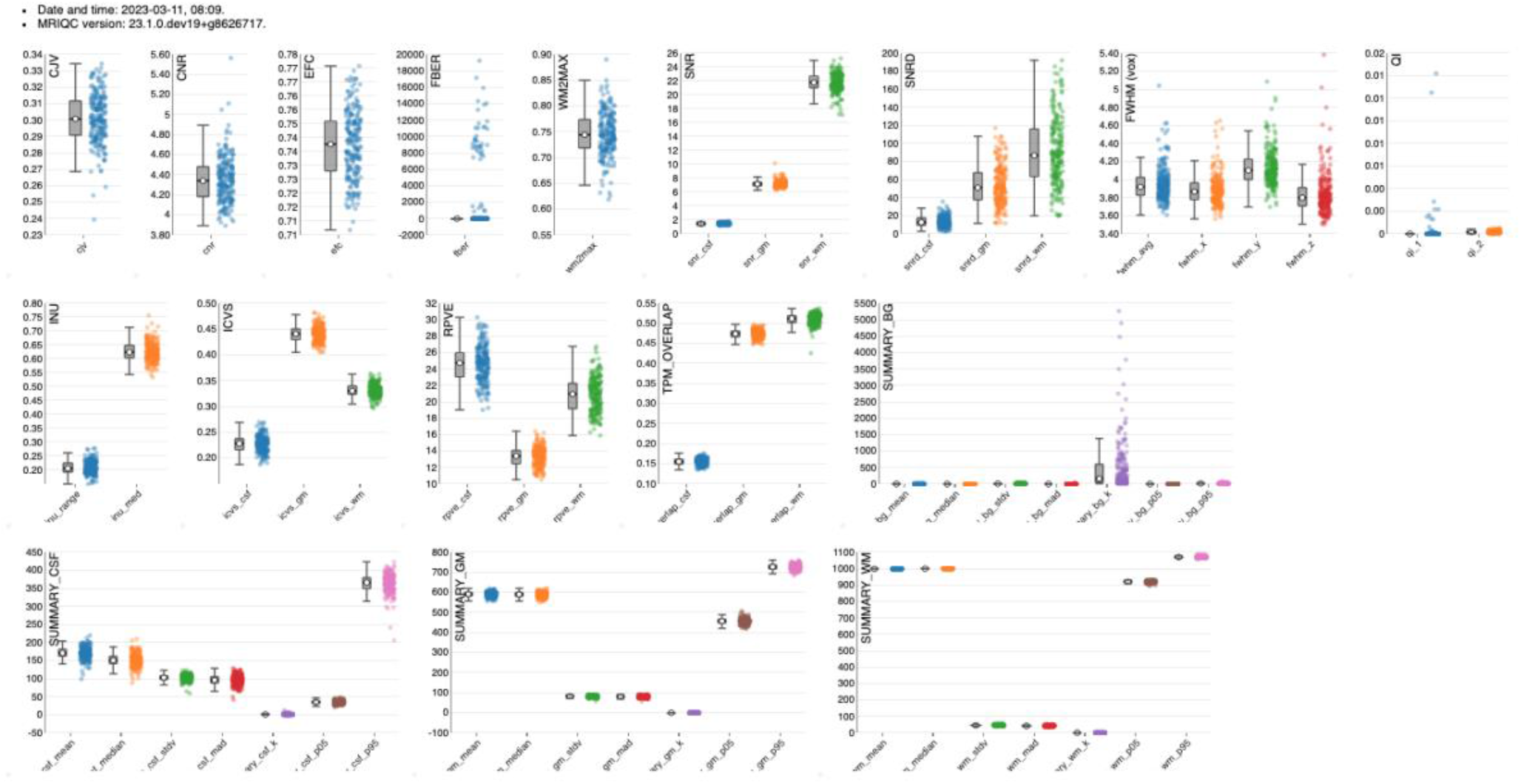
MRIQC: group T1w report. The group report for each MRI modality shows the distribution of each IQM across participants.

#### 3.2 Visual inspection of group reports (⬤TIMING 10-15 min)

The group reports can be used to investigate the range and clustering of each IQM, as well as identify participants that are outliers in some IQM distribution. Individual reports can be viewed by clicking on data points, so that researchers can see how the visual reports are impacted by high, low, and average IQM values.

### 4 Post assessment of IQMs and ratings

#### 4.1 QC results management (⬤TIMING 10-15 min)

After all scans have been inspected, ratings and notes can be exported from *Q’kay*, or collated from downloaded rating JSONs. If more than one rater inspected reports, inter-rater reliability can be assessed, and discrepancies (especially on scans flagged to be excluded) should be noted and resolved. Ratings and exclusion status can be added to the BIDS-required `participants.tsv` file to select scans to feed into downstream preprocessing tools (like *fMRIPREP)* or incorporation into analytic workflows.

▲**CRITICAL** In general, “unuseable” data should not be deleted in case it may be suitable for another analysis, or with future preprocessing advances. Sharing “unuseable” data also supports the development and improvement of automatic QC algorithms. *MRIQC* reports, rating JSONs, and SOPs detailing exclusion criteria should be stored with the raw data to facilitate data sharing and reuse.

#### 4.2 Automated classification of images (optional)

For large datasets, automated QC may be desired to decrease the researcher time required to manually rate images. This can be achieved by training a classifier to predict ratings from the IQMs, using a set of labeled ratings as the training data. Previous experiments using IQMs to predict image quality found that these models can be influenced by the sensitivity of IQMs to scan parameters and site, and may not generalize from one dataset/site to another^7^. More details about the IQMs and how to create a classifier can be found in the *NiPreps* ISMRM QC book^u^.

## Troubleshooting

Some of the most common pitfalls encountered by *MRIQC* users relate to resource management and other set-up settings (step 1.3 in <u>Data acquisition</u> or step 2.1 in <u>Execute *MRIQC*</u> subsection), as suggested by the many questions the source code repository^v^ and the NeuroStars.org channel each receive weekly. In particular, the limitations imposed by each HPC system, and the particularities of the *Apptainer* container framework may require some troubleshooting.

### Invalid BIDS dataset

A fairly common reason for *MRIQC* to fail is the attempt to use non-BIDS data. Therefore, the first troubleshooting step is running the *BIDS-Validator*. When using containers to run *MRIQC*, if the container does not have access to the data, the validator will flag the dataset as invalid. Containers are a confined computation environment and they are not allowed to access the host’s filesystems, unless explicit measures are taken to grant access (i.e., mounting or binding filesystems using the `-v` flag for *Docker* or the equivalent --bind for *Apptainer*). Therefore, when using containers with a valid BIDS dataset, the “invalid BIDS dataset” could be a symptom of failure to access the data from the host.

### Network filesystem errors

*MRIQC* is built on *Nipype*^24^, a neuroimaging workflow framework that uses the filesystem to coordinate the data flow during execution. Network filesystems may exhibit large latencies and temporary inconsistencies that may break execution. Setting the “working directory” option to a local, synchronized filesystem will preempt these issues.

### Memory errors

When running on systems with restrictive memory overcommit policies (frequently found in multi-tenant HPC systems), the *MRIQC* virtual memory footprint may become too large, and the process will be stopped by the job scheduler or the operating system kernel. The recommendation in this scenario is to split (parallelize) processing across subjects (Box 2 showcases a solution). Alternatively, when running on a system with 8GB RAM or less, *MRIQC* is likely to exceed physical memory limits. This scenario is particularly common when running the container version of *MRIQC*, because the container has access to a very low physical memory allocation. For example, *Docker* typically limits memory to 2GB by default on *macOS* and *Windows* systems. In this case, the solution is to increase the memory allocation available to *MRIQC* (via adequate settings of the container engine and/or upgrading the hardware).

### Hard disk quotas

Shared systems generally limit the hard disk space a user can use. Please allocate enough space for both intermediate and final results. Remove intermediate results as soon as satisfied with the final results to free up scratch space.

### PyBIDS indexing

For large datasets, *PyBIDS* indexing inflates run time. To optimize run time, a pre-indexed cache of the BIDS structure and metadata can be created with *PyBIDS*, and will substantially speed *MRIQC* up:

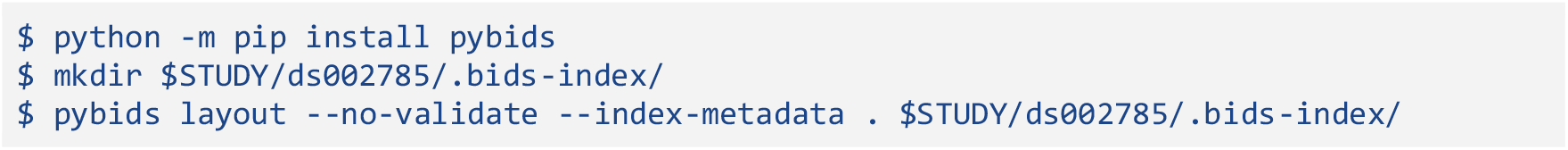

Once the pre-indexed cache is built, you should inform *MRIQC* of its location by setting the command line argument `--bids-database-dir $STUDY/ds002785/.bids-index/`.

### NeuroStars forum

Many other frequently asked questions are found and responded to at NeuroStars.org. New support requests are welcome via this platform.

## Anticipated results

The successful application of this protocol produces the following outcomes: visual reports and qualitative ratings from assessing those reports, and quantitative metrics of image quality for each scan in the dataset. These can be used to prepare the dataset for analysis by identifying scans that do or do not meet the exclusion criteria. Additionally, this information can be used to generate data to train a classifier for your scanner, enabling automated image quality assessment.

## Data availability

The MRI data used in this protocol are publicly available at OpenNeuro.org (doi: 10.18112/openneuro.ds002785.v2.0.0). *MRIQC* reports generated as part of this protocol are available at https://mriqc.s3.amazonaws.com/index.html#Hagen2024/.

## Acknowledgements

We thank the invaluable feedback on this report from Jo Etzel, Ph.D., as well as her continued support to our QA/QC endeavors. This work was supported by Chan-Zuckerberg-Initiative (EOSS5-266, O.E.), the NIMH (R24MH114705 and R24MH117179, R.A.P.; RF1MH121867, O.E., A.R., and R.A.P.). C.P. and O.E. receive support from the Swiss National Science Foundation —SNSF— (grant #185872, O.E.). This material is based upon work supported by the U.S. Department of Energy, Office of Science, Office of Advanced Scientific Computing Research, Department of Energy Computational Science Graduate Fellowship under Award Number DE-SC0023112 (M.P.H). This work was supported in part by the NIH National Institute on Mental Health (T32MH122394) and National Institute on Aging (F31AG062067, S.S.). MN was supported by the BRAIN initiative (MH002977-01).

## Disclaimer

This report was prepared as an account of work sponsored by an agency of the United States Government. Neither the United States Government nor any agency thereof, nor any of their employees, makes any warranty, express or implied, or assumes any legal liability or responsibility for the accuracy, completeness, or usefulness of any information, apparatus, product, or process disclosed, or represents that its use would not infringe privately owned rights. Reference herein to any specific commercial product, process, or service by trade name, trademark, manufacturer, or otherwise does not necessarily constitute or imply its endorsement, recommendation, or favoring by the United States Government or any agency thereof. The views and opinions of authors expressed herein do not necessarily state or reflect those of the United States Government or any agency thereof.

This content is solely the responsibility of the authors and does not necessarily represent the official views of the National Institutes of Health.

## Author contribution statement

MPH, CP, AR and OE contributed formal analysis, investigation, methodology, validation. MPH, CP and OE wrote the original draft. MPH, CP, and JKL developed and validated the protocol. PB, RAP, AR, and OE contributed to conceptualization, interpretation, overall framing of the protocol, funding acquisition, project administration, resources, and supervision. EM, JHL, MN, MG, SS and TG contributed to software, methodology and validation. All the authors have contributed software and/or documentation, read the manuscript, and edited/revised the original draft and previous versions.

## Footnotes

a https://mriqc.readthedocs.io/en/latest/#

b https://www.nipreps.org/

c https://qsiprep.readthedocs.io/en/latest/preprocessing.html#visual-reports

d www.nipreps.org/dmriprep-viewer/

e https://fibr.dev/

f https://github.com/nipreps/qkay

g www.github.com

h www.nipreps.org/sops-cookiecutter/

i www.frontiersin.org/articles/10.3389/fnimg.2022.1073734/full#supplementary-material

j www.axonlab.org/hcph-sops/data-collection/participant-prep/

k https://bids-standard.github.io/bids-validator/

l https://github.com/bids-standard/bids-validator?tab=readme-ov-file#quickstart

m https://mriqc.readthedocs.io/en/stable/docker.html

n https://handbook.datalad.org/en/latest/

° https://mriqc.readthedocs.io/en/stable/

p https://mriqc.s3.amazonaws.com/index.html#aomic-piop1/mriqc-23.0.0rc0/

q https://github.com/nipreps/mriqc/tree/master/docs/source/resources

r https://neurostars.org/

s https://github.com/nipreps/mriqc/issues?q=is%3Aissue

t https://mriqc.readthedocs.io/en/stable/measures.html

u https://www.nipreps.org/qc-book/

v https://github.com/nipreps/mriqc/issues?q=is%3Aissue

## Notes

### Competing Interest Statement

The authors have declared no competing interest.

## References

1. Snoek, L. et al. The Amsterdam Open MRI Collection, a set of multimodal MRI datasets for individual difference analyses. Sci. Data 8, 85 (2021).

2. Power, J. D., Barnes, K. A., Snyder, A. Z., Schlaggar, B. L. & Petersen, S. E. Spurious but systematic correlations in functional connectivity MRI networks arise from subject motion. NeuroImage 59, 2142–2154 (2012).

3. Yendiki, A., Koldewyn, K., Kakunoori, S., Kanwisher, N. & Fischl, B. Spurious group differences due to head motion in a diffusion MRI study. NeuroImage 88, 79–90 (2014).

4. Reuter, M. et al. Head motion during MRI acquisition reduces gray matter volume and thickness estimates. NeuroImage 107, 107–115 (2015).

5. Alexander-Bloch, A. et al. Subtle in-scanner motion biases automated measurement of brain anatomy from in vivo MRI. Hum. Brain Mapp. 37, 2385–2397 (2016).

6. Etzel, J. A. Efficient evaluation of the Open QC task fMRI dataset. Front. Neuroimaging 2, (2023).

7. Esteban, O. et al. MRIQC: Advancing the automatic prediction of image quality in MRI from unseen sites. PLOS ONE 12, e0184661 (2017).

8. Esteban, O. et al. Crowdsourced MRI quality metrics and expert quality annotations for training of humans and machines. Sci. Data 6, 1–7 (2019).

9. Lepping, R. J. et al. Quality control in resting-state fMRI: the benefits of visual inspection. Front. Neurosci. 17, (2023).

10. Provins, C., MacNicol, E., Seeley, S. H., Hagmann, P. & Esteban, O. Quality control in functional MRI studies with MRIQC and fMRIPrep. Front. Neuroimaging 1, (2023).

11. Gardner, E. A. et al. Detection of degradation of magnetic resonance (MR) images: Comparison of an automated MR image-quality analysis system with trained human observers. Acad. Radiol. 2, 277–281 (1995).

12. Woodard, J. P. & Carley-Spencer, M. P. No-Reference image quality metrics for structural MRI. Neuroinformatics 4, 243–262 (2006).

13. Gedamu, E. L., Collins, D. l. & Arnold, D. L. Automated quality control of brain MR images. J. Magn. Reson. Imaging 28, 308–319 (2008).

14. Mortamet, B. et al. Automatic quality assessment in structural brain magnetic resonance imaging. Magn. Reson. Med. 62, 365–372 (2009).

15. Shehzad, Z. et al. The Preprocessed Connectomes Project Quality Assessment Protocol - a resource for measuring the quality of MRI data. in INCF Neuroinformatics (Front Neurosci, Cairns, Australia, 2015). doi:10.3389/conf.fnins.2015.91.00047.

16. Pizarro, R. A. et al. Automated Quality Assessment of Structural Magnetic Resonance Brain Images Based on a Supervised Machine Learning Algorithm. Front. Neuroinformatics 10, (2016).

17. Backhausen, L. L. et al. Quality Control of Structural MRI Images Applied Using FreeSurfer—A Hands-On Workflow to Rate Motion Artifacts. Front. Neurosci. 10, (2016).

18. Alfaro-Almagro, F. et al. Image processing and Quality Control for the first 10,000 brain imaging datasets from UK Biobank. NeuroImage (2017) doi:10.1016/j.neuroimage.2017.10.034.

19. White, T. et al. Automated quality assessment of structural magnetic resonance images in children: Comparison with visual inspection and surface-based reconstruction. Hum. Brain Mapp. 39, 1218–1231 (2018).

20. Keshavan, A., Yeatman, J. & Rokem, A. Combining citizen science and deep learning to amplify expertise in neuroimaging. bioRxiv 363382 (2018) doi:10.1101/363382.

21. Klapwijk, E. T., van de Kamp, F., van der Meulen, M., Peters, S. & Wierenga, L. M. Qoala-T: A supervised-learning tool for quality control of FreeSurfer segmented MRI data. NeuroImage 189, 116–129 (2019).

22. Esteban, O. et al. MRIQC: Advancing the automatic prediction of image quality in MRI from unseen sites. PLOS ONE 12, e0184661 (2017).

23. Gorgolewski, K. J. et al. The brain imaging data structure, a format for organizing and describing outputs of neuroimaging experiments. Sci. Data 3, 160044 (2016).

24. Gorgolewski, K. et al. Nipype: a flexible, lightweight and extensible neuroimaging data processing framework in Python. Front. Neuroinformatics 5, 13 (2011).

25. Esteban, O. et al. Analysis of task-based functional MRI data preprocessed with fMRIPrep. Nat. Protoc. 15, 2186–2202 (2020).

26. Poldrack, R. A. et al. The Past, Present, and Future of the Brain Imaging Data Structure (BIDS). ArXiv 2309.05768v2 (2024).

27. MacNicol, E. E. et al. Extending MRIQC to rodents: image quality metrics for rat MRI. in Annual Meeting of the European Society for Molecular Imaging (EMIM) vol. 17 PW23-913 (Thessaloniki, Greece, 2022).

28. Provins, C. et al. Defacing biases in manual and automatic quality assessments of structural MRI with MRIQC. Peer Community Regist. Rep. Regist. Rep. Consid. Stage 1.

29. Provins, C. et al. Defacing biases manual and automated quality assessments of structural MRI with MRIQC. Preprint at 10.31219/osf.io/t9ehk (2022).

30. Marcus, D. S. et al. Human Connectome Project informatics: Quality control, database services, and data visualization. NeuroImage 80, 202–219 (2013).

31. Keshavan, A., Yeatman, J. D. & Rokem, A. Combining Citizen Science and Deep Learning to Amplify Expertise in Neuroimaging. Front. Neuroinformatics 13, 29 (2019).

32. Richie-Halford, A. et al. An analysis-ready and quality controlled resource for pediatric brain white-matter research. Sci. Data 9, 616 (2022).

33. Garcia, M., Dosenbach, N. & Kelly, C. BrainQCNet: a Deep Learning attention-based model for the automated detection of artifacts in brain structural MRI scans. Imaging Neurosci. (2024) doi:10.1162/imag_a_00300.

34. Sanchez, T. et al. FetMRQC: A robust quality control system for multi-centric fetal brain MRI. Med. Image Anal. 97, 103282 (2024).

35. Heunis, S. et al. Quality and denoising in real-time functional magnetic resonance imaging neurofeedback: A methods review. Hum. Brain Mapp. 41, 3439–3467 (2020).

36. Cox, R. W. & Hyde, J. S. Software tools for analysis and visualization of fMRI data. NMR Biomed. 10, 171–178 (1997).

37. Koush, Y. et al. OpenNFT: An open-source Python/Matlab framework for real-time fMRI neurofeedback training based on activity, connectivity and multivariate pattern analysis. NeuroImage 156, 489–503 (2017).

38. Dosenbach, N. U. F. et al. Real-time motion analytics during brain MRI improve data quality and reduce costs. NeuroImage 161, 80–93 (2017).

39. Keshavan, A. et al. Mindcontrol: A web application for brain segmentation quality control. NeuroImage 170, 365–372 (2018).

40. Rosen, A. F. G. et al. Quantitative assessment of structural image quality. NeuroImage 169, 407–418 (2018).

41. Niso, G. et al. Open and reproducible neuroimaging: from study inception to publication. NeuroImage 119623 (2022) doi:10.1016/j.neuroimage.2022.119623.

42. Halchenko, Y. O. et al. Open Brain Consent: make open data sharing a no-brainer for ethics committees. Zenodo (2018) doi:10.5281/zenodo.1411525.

43. Markiewicz, C. J. et al. The OpenNeuro resource for sharing of neuroscience data. eLife 10, e71774 (2021).

44. Avants, B. B. et al. A reproducible evaluation of ANTs similarity metric performance in brain image registration. NeuroImage 54, 2033–44 (2011).

45. Jenkinson, M., Beckmann, C. F., Behrens, T. E. J., Woolrich, M. W. & Smith, S. M. FSL. NeuroImage 62, 782–790 (2012).

46. Abraham, A. et al. Machine learning for neuroimaging with scikit-learn. Front. Neuroinformatics 8, (2014).

47. Savary, E., Provins, C., Sanchez, T. & Esteban, O. Q’kay: a manager for the quality assessment of large neuroimaging studies. in (29th Annual Meeting of the Organization for Human Brain Mapping (OHBM), 2023). doi:10.31219/osf.io/edx6t.

48. Hollmann, S. et al. Ten simple rules on how to write a standard operating procedure. PLoS Comput. Biol. 16, e1008095 (2020).

49. OHBM2018 - ReproIn. Google Docs https://docs.google.com/document/d/1EsQH3kVTdIstvjNoB-9vJxB8kGhtG4guKNaPVe-EP54/edit?usp=embed_facebook.

50. Li, X., Morgan, P. S., Ashburner, J., Smith, J. & Rorden, C. The first step for neuroimaging data analysis: DICOM to NIfTI conversion. J. Neurosci. Methods 264, 47–56 (2016).

51. Halchenko, Y. O. et al. Open Source Software: Heudiconv. Zenodo (2018) doi:10.5281/zenodo.1306159.

52. Covitz, S. et al. Curation of BIDS (CuBIDS): A workflow and software package for streamlining reproducible curation of large BIDS datasets. NeuroImage 263, 119609 (2022).

53. Schwarz, C. G. et al. Changing the face of neuroimaging research: Comparing a new MRI de-facing technique with popular alternatives. NeuroImage 231, 117845 (2021).

54. White, T., Blok, E. & Calhoun, V. D. Data sharing and privacy issues in neuroimaging research: Opportunities, obstacles, challenges, and monsters under the bed. Hum. Brain Mapp. 43, 278–291 (2022).

55. Yoo, A. B., Jette, M. A. & Grondona, M. SLURM: Simple Linux Utility for Resource Management. in Job Scheduling Strategies for Parallel Processing (eds. Feitelson, D., Rudolph, L. & Schwiegelshohn, U.) 44–60 (Springer Berlin Heidelberg, Seattle, WA, USA, 2003). doi:10.1007/10968987_3.

56. Kurtzer, G. M., Sochat, V. & Bauer, M. W. Singularity: Scientific containers for mobility of compute. PLOS ONE 12, e0177459 (2017).

57. Gorgolewski, K. J. et al. BIDS Apps: Improving ease of use, accessibility, and reproducibility of neuroimaging data analysis methods. PLOS Comput. Biol. 13, e1005209 (2017).

58. Esteban, O. et al. fMRIPrep: a robust preprocessing pipeline for functional MRI. Nat. Methods 16, 111–116 (2019).

59. Power, J. D. A simple but useful way to assess fMRI scan qualities. NeuroImage 154, 150–158 (2017).

60. Patriat, R., Molloy, E., Birn, R., Guitchev, T. & Popov, A. Using Edge Voxel Information to Improve Motion Regression for rs-fMRI Connectivity Studies. Brain Connect. 5, 582–595 (2015).

61. Gudbjartsson, H. & Patz, S. The rician distribution of noisy mri data. Magn. Reson. Med. 34, 910–914 (1995).

